# In the absence of reproductive isolation – Extensive gene flow after speciation

**DOI:** 10.1101/622019

**Authors:** Xinfeng Wang, Zixiao Guo, Ziwen He, Shaohua Xu, Shao Shao, Sen Li, Ming Yang, Qipian Chen, Cairong Zhong, Zhongyi Wu, Norman C. Duke, Suhua Shi

## Abstract

In the conventional view, species are separate gene pools delineated by reproductive isolation (RI). However, species may also be delineated by merely a small set of “speciation genes” without full RI. It is thus important to know whether “good species” (defined by the “secondary sympatry” test) do continue to exchange genes. Here, we carry out sequencing and *de novo* high-quality assembly of the genomes of two closely related mangrove species (*Rhizophora mucronata* and *R. stylosa*). Whole-genome re-sequencing of individuals across their range on the tropical coasts shows their genomes to be well delineated in allopatry. They became sympatric in northeastern Australia but remain distinct species in contact. Nevertheless, their genomes harbor ∼ 4,000 to 10,000 introgression blocks, each averaging only about 3-4 Kb. These fine-grained introgressions indicate that gene flow has continued long after speciation. Non-introgressable “genomic islets,” averaging only 1.4 Kb, may contribute to speciation as they often harbor diverging genes underlying flower development and gamete production. In conclusion, RI needs not be the main criterion of species delineation even though all species would eventually be fully reproductively isolated.

## Introduction

Biological species are generally defined as taxa that do not exchange genes due to various forms of reproductive isolation (RI). RI mechanisms include ecological, behavioral, and reproductive incompatibilities (*1, 2*). These mechanisms are the foundation of the Biological Species Concept (BSC; (*3*–*5*)) and have been accepted as both necessary and sufficient for species to evolve along diverging paths.

Strictly speaking, two true species should be separated by a combination of RI mechanisms such that they cease to exchange genes anywhere in their genomes. RI needs to be complete to avoid the logical quagmire of defining how much isolation is enough. In addition, RI assumes a very high degree of genetic cohesiveness within each species such that any exchange would be harmful (*3, 5, 6*). Complete RI, however, is difficult to ascertain. For example, many good species appear to be only partially isolated as they produce sterile hybrids in one sex but the other sex is highly fertile (*7*–*11*). In many other cases, there is no evidence even for partial RI (*12*–*15*).

Against this backdrop, Wu (*5*) questioned the basic assumptions of BSC in asserting full RI across the whole genome. In reality, only a small fraction of the genome may be responsible for differentiation between species in morphology, behavior, reproduction, and ecology (i.e., “speciation genes”). In this genic view of speciation (GVS), genome regions not germane to speciation should be easily interchangeable between species. In short, BSC asserts that true species should be fully reproductively isolated. The alternative GVS, by allowing species to extensively share genetic material, would abandon RI as the defining concept of species. The central question is, therefore, “do true species extensively exchange genes?”

### Conditions for detecting post-speciation gene flow

To answer the question above, we may use the model outlined in Fig. 1A-1D to evaluate the conditions for detecting post-speciation gene flow. First, the species status needs to be defined by unambiguous criteria. We suggest the “secondary sympatry” test to delineate species whereby two populations have evolved into putatively full species before coming into secondary contact. In sympatry, true species should be able to maintain their biological distinction over an extended period, even with ample opportunities for genetic exchanges to merge into a hybrid swarm (Fig. 1A). The sympatry test is important as seen in previous debates on this issue (*6, 16*–*18*). In many cases, the species may not pass the sympatry test as shown in Fig. 1B (see the legends). Furthermore, while the extensive genomic literature has convincingly demonstrated gene flow during speciation (*6*), the inferred gene flow most likely happened in the early stages (*6*). Note that the focus here is on post-speciation gene flow.

**Figure 1.**
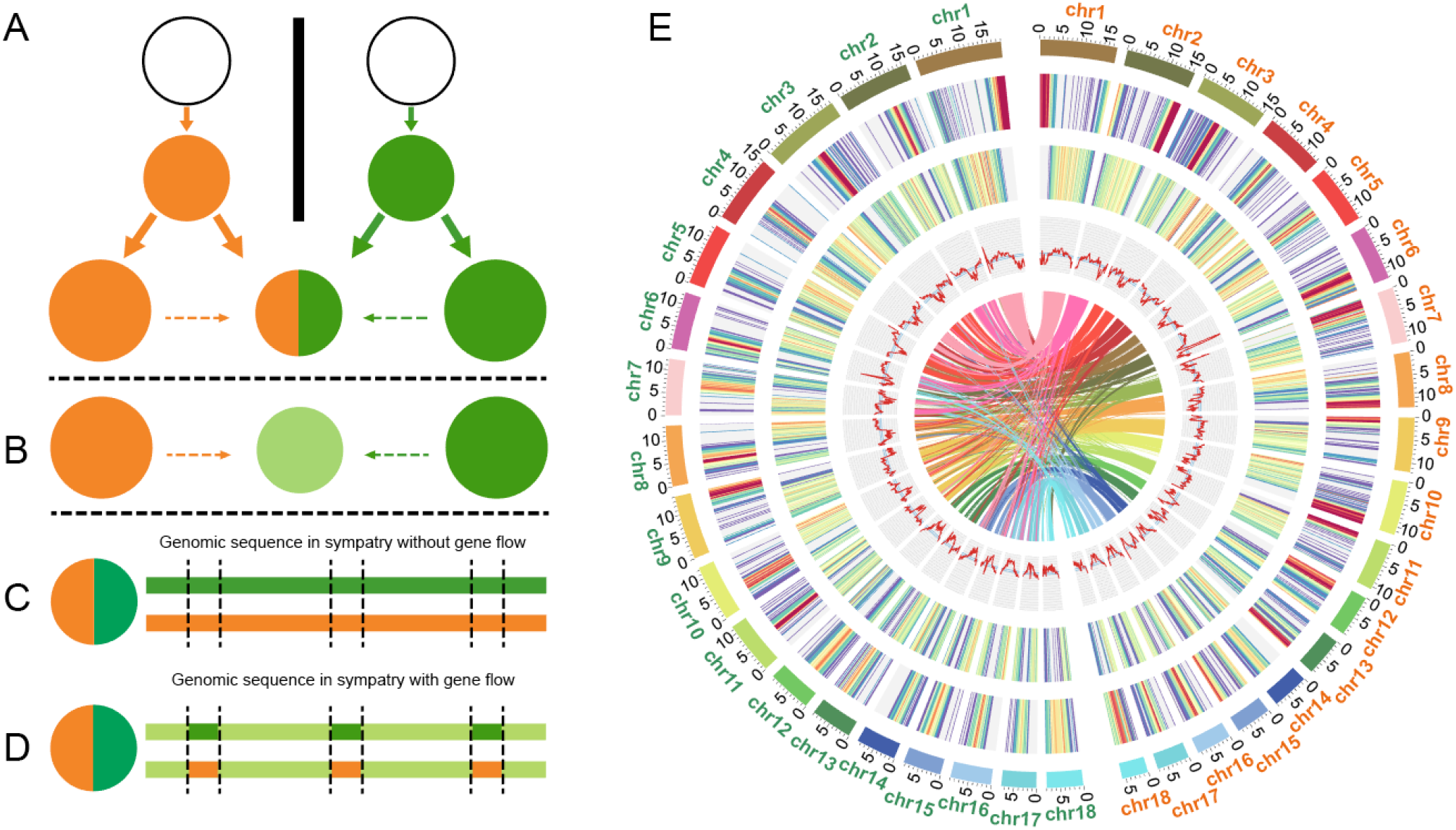
Models of post-speciation gene flow and the genomic structure in such models. (A) Two populations undergo allopatric speciation, resulting in two good species depicted in orange and green, respectively. In an area away from the center of each species’ distribution, the two putative species re-establish sympatry. In sympatry, the two species retain their biological characters (shown in their distinctive colors), passing the “secondary sympatry” test. (B) In many cases, the two putative species hybridize and fuse (indicated by the olive color) in sympatry. They are not considered true species. (C-D) When the two species pass the sympatry test, there are two possibilities. In (C), the two species do not exchange genes at all. This is the most common scenario as observed in many secondary sympatry cases. (In such cases, the hybrid zones may show some admixture although the hybrids are usually quickly eliminated by natural selection.) (D) The true case of post-speciation gene flow. The genomic sequences of the two species show signs of genetic exchanges in the genomic segments of olive color. The small genomic segments flanked by dotted lines are non-introgressable and may harbor “speciation genes.” It should be noted that the patterns of introgression are transient. As the two species continue to diverge, gene flow within the same species will homogenize the genomic sequences of each species, thus reversing the genomic patterns back to that shown in (C). (E) The genomes of *R. mucronata* and *R. stylosa*. Circular tracks represent, from outer to inner, top 18 longest scaffolds of *R. mucronata* (chr1-18 in orange) and *R. stylosa* (chr1-18 in green), percentage of repeats (1–99%), gene density (0–49), GC content (29.61–51.97%) and the spectrum of inter-specific collinear analysis (each line connects one pair of homologous genes and a cluster of such lines represents one collinear block). All statistics are calculated for windows of 200 Kb.

Once good species are confirmed by the test, it would be possible to check for past genetic exchanges discernible in their genomes. Importantly, Wang et al.’s (*6*) extensive survey finds no convincing evidence for post-speciation gene flow. Such evidence may not be readily accessible as the window of time for convincing observations is quite narrow. If it is too early, the species may not pass the “secondary sympatry” test (Fig. 1B). On the other hand, the footprint of post-speciation gene flow may not last long as subsequent divergence and gene flow would erase it (Fig. 1C).

The true signature of post-speciation gene flow requires two signals. First, allopatric populations of these species should show the pattern depicted in Fig. 1C. Second, sympatric populations would show the Fig. 1D pattern. The contrast between the two signals ensures that the gene flow in sympatry has indeed happened after speciation has completed. As stated above, gene flow during speciation, as concluded in many studies (*6, 19, 20*), most likely happens in the early stages. The model illustrated in Fig. 1A-1D shows the challenges of proving post-speciation gene flow. A convincing proof, requiring careful experimental design to identify species at the right stage and in the right place of evolution, would be highly conceptually significant. This study provides such a proof.

## Results

Two closely related mangrove species – *Rhizophora mucronata* vs. *R. stylosa* (*21*–*28*) appear most suited to answer the central question we want to address. Mangroves are woody plants that have colonized intertidal zones of tropical coasts (*21*–*23, 29*). Because of the narrow band of suitable habitats along the coasts (or near river mouths), global mangrove distributions are essentially one-dimensional, making them ideal for biogeographical studies of speciation. We shall begin the analyses by first presenting the genomes of the two mangrove species.

### De novo assembly R. mucronata and R. stylosa genomes

We sequenced the genome of one *R. mucronata* and one *R. stylosa* individual, with estimated genome size of ∼300 Mb and ∼370 Mb, respectively (Supplementary Table S1; see Supplementary Table S2 for details). Although the genomes of these species have been re-sequenced before (*22*), the aims of this study demand accurate assembly of their own genomes in order to assess possible fine-grained introgressions. We therefore carried out de-novo sequencing and attained the chromosome-level assembly in this study. Re-sequencing of individuals from their geographical ranges is based on this new assembly.

We obtained 33.84 Gb (gigabases) and 36.13 Gb of raw data, corresponding to 142X coverage of the assembled genomes (Supplementary Table S2). We assembled 237.85 Mb (megabases) of *R. mucronata* sequences into 14,496 scaffolds with N50 at 12.03 Mb and 18 scaffolds (chr1-18) >= 5 Mb. The assembled chr1-18 account for 84.38% (200.70 Mb) of the genome and correspond to the diploid chromosome number (2n = 36, Fig. 1E, Supplementary Table S1, and Fig. S1). The assembled *R. stylosa* genome is 253.54 Mb, containing 9,750 scaffolds with N50 at 12.59 Mb. The top 18 scaffolds account for 219.66 Mb, or 86.63%, of the genome (Fig. 1E, Supplementary Table S1, and Fig. S2). The estimated *R. mucronta* genome coverage (94.2%) and mapping rate (94.52%) are both high. The numbers are 92.74% and 94.3% for *R. stylosa* (Supplementary Tables S1-S2). These results indicate high-quality de novo genomes approaching chromosome-level completeness. A total of 26,540 protein-coding genes were annotated and classified into 19,089 families in the *R. mucronata* genome, while 30,375 genes from 19,018 families were identified in *R. stylosa* (Supplementary Table S1).

To examine the evolution of *R. mucronata* and *R. stylosa* genome structure in the larger phylogenetic context, we compare their genomes to those of *Carallia longipes* (unpublished data), *Bruguiera gymnorrhiza* (*29*) and *Rhizophora apiculate* (*22*). *C. longipes* is from one of the closest non-mangrove genera in the Rhizophoraceae family. 20,971 gene families were identified among the five species, with 6,014 single-copy. Using these single-copy orthologs, we reconstructed the species phylogeny and estimated divergence times (Supplementary Fig. S3). The phylogeny shows that *R. mucronata* and *R. stylosa* are the most closely related pair diverging ∼3.75 Mya (with 95% confidence interval [3.15, 4.35] Mya; Supplementary Table S3 and Fig. S3). The inter-specific Ks (∼0.0031) and the genomic divergence (*D*_*xy*_ = 0.0031) all support the close relationship between *R. mucronata* and *R. stylosa* (Supplementary Table S2 and Fig. S4). We identify 663 collinear blocks between the two species that harbor 18,705 genes in *R. mucronata* and 18,952 in *R. stylosa* (Fig. 1E).

### R. mucronata and R. stylosa genomic diversity

*Rhizophora mucronata* is widely distributed in the Indo-Western Pacific (IWP) region, particularly to the west of the Strait of Malacca and all the way to East Africa. In contrast, *R. stylosa* extends eastward from the Strait of Malacca to western Pacific Islands (Fig. 2A). The two species have been reported to overlap in scattered locales along several western Pacific coastlines. However, in our own field trips, their relative abundance is often skewed in favor of one species and their co-occurrence has been rarely found. The sole exception in our collection is in the Daintree River (DR) area of northeastern Australia, where both species are quite abundant (Fig. 2A).

**Figure 2.**
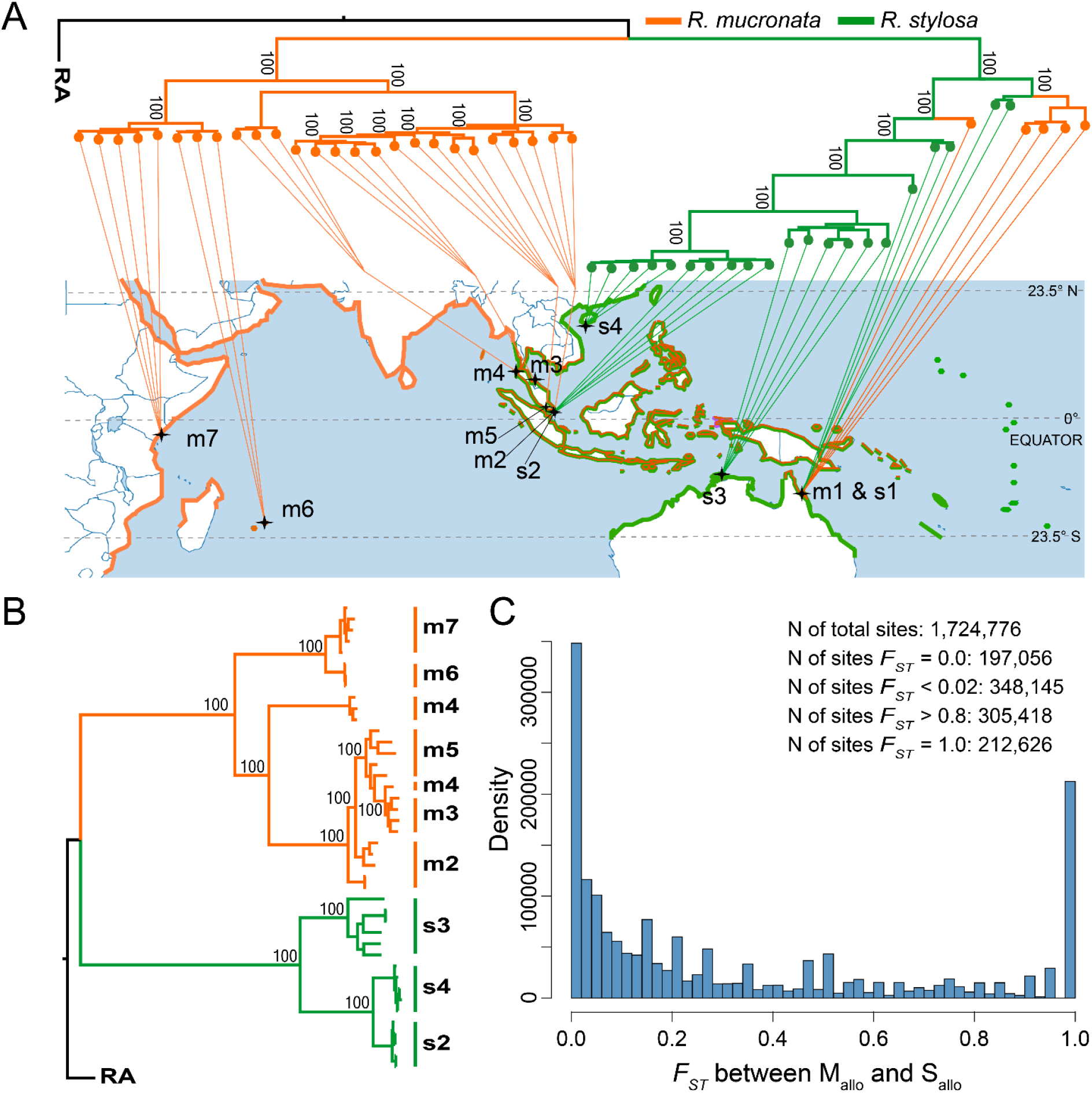
Phylogenetic relationships and biogeographical observations. (A) The biogeography and genealogy of *R. mucronata* and *R. stylosa*. The maximum-likelihood (ML) tree, generated by RAxML with 100 bootstraps, is superimposed on the biogeography. The numbers on the nodes indicate support values. *R. mucronata* is colored in orange while *R. stylosa* is in green. (B) The same phylogeny excluding the sympatric m1 and s1 samples from the Daintree area. We denote the allopatric populations as M_allo_ (m2-m7) and S_allo_ (s2-s4). (C) The spectrum of the *F*_*ST*_ statistic between the M_allo_ and S_allo_ samples.

We collected 21 *R. stylosa* individuals from four locations (labeled s1-s4) and 31 *R. mucronata* samples from seven locations (m1-m7) for population genomic studies (Fig. 2A and Supplementary Table S4). Note that m1 and s1 refer to the sympatric DR samples. Whole genomes of all samples were sequenced on the Illumina Hiseq 2000 platform, yielding a mean depth of 16X (ranging from 12X to 22X) (Supplementary Table S5 and Table S6). Short reads from each individual were mapped to the de novo *R. mucronata* genome, with average genomic coverage of 81% (80%-83%, Supplementary Table S4). The level of genetic diversity shows two patterns. Low genetic diversity is found in all allopatric populations (average **θ**_π_ at 0.62 and 0.60 per Kb for *R. mucronata* m2-m7 and *R. stylosa* s2-s4, respectively, see also Supplementary Tables S2 and S4). The level is much higher in the sympatric DR populations (**θ**_π_ = 1.37/Kb and 2.09/Kb, respectively). Watterson’s estimates (**θ**_w_) are similar (Supplementary Table S4, see Materials and Methods).

### Divergence between the two species in allopatry

Genomic divergence (*D*_*xy*_) between the two species is 4.14 × 10^−3^ per site (Supplementary Table S7). We first constructed a Maximum Likelihood (ML) tree using RAXML (*30*) on 31 *R. mucronata* and 21 *R. stylosa* individual sequences from the 11 populations. The ML tree bifurcates with a clear delineation between species across all allopatric populations. However, the m1 and s1 (i.e., DR) samples show strong signs of admixture as they are “in the middle” of the bifurcated tree (Fig. 2A). When the DR samples are removed, the phylogeny shows clear delineation (Fig. 2B). These two trees are robust when rebuilt using the ML method in IQTREE (*31*) or the Neighbor-Joining (NJ) method in MEGA7 (*32*) (Supplementary Fig. S5 and S6). The monophyletic delineation of *R. mucronata* and *R. stylosa* in allopatry is also supported by a principal component analysis (PCA (*33*), Supplementary Fig. S7).

We detected 1.7 million variable sites across all populations of the two species (Supplementary Table S8). We first partition these sites by excluding the DR samples (see Materials and Methods). Each site is then represented by an *F*_*ST*_ value with *F*_*ST*_ = 0 indicating no differentiation between the two species in allopatry and *F*_*ST*_ = 1 indicating complete differentiation. Figure 2C shows the U-shaped distribution where the abundance of sites at the far right reveals the extensive differentiation between species. Such a U-shape distribution is typical of species diverging with little gene flow (*34*).

The two species can be easily distinguished in the field. *R. mucronata* tends to settle in the less saline and further upstream habitats in comparison with *R. stylosa* found in saline habitats closer to river mouths (see Fig. 3E). The two species also differ substantially in overall tree morphology (see Fig. 3D). They are most readily distinguished by the reproductive characters of the flower, in particular the style length (*23, 24*) as pictured in Fig. 3A. The morphological differences between *R. mucronata* and *R. stylosa* across populations are shown in Fig. 3C. *R. mucronata* is readily distinguished by its short style, between 0.9 and 1.6mm (Fig. 3A). In contrast, the *R. stylosa* style is long, 2.4-5.3 mm (Fig. 3A), with no overlap between the two species (Fig. 3C) (*23, 24*). While the style length varies from locale to locale in both species, this trait is species-diagnostic across locales. Additional, albeit less stable, diagnostic morphological characters are listed in the Supplement (Table S9). In short, we show that *R. mucronata* and *R. stylosa* have diverged in their genomes, geographical distribution, habitat choice, and various morphological characters.

**Figure 3.**
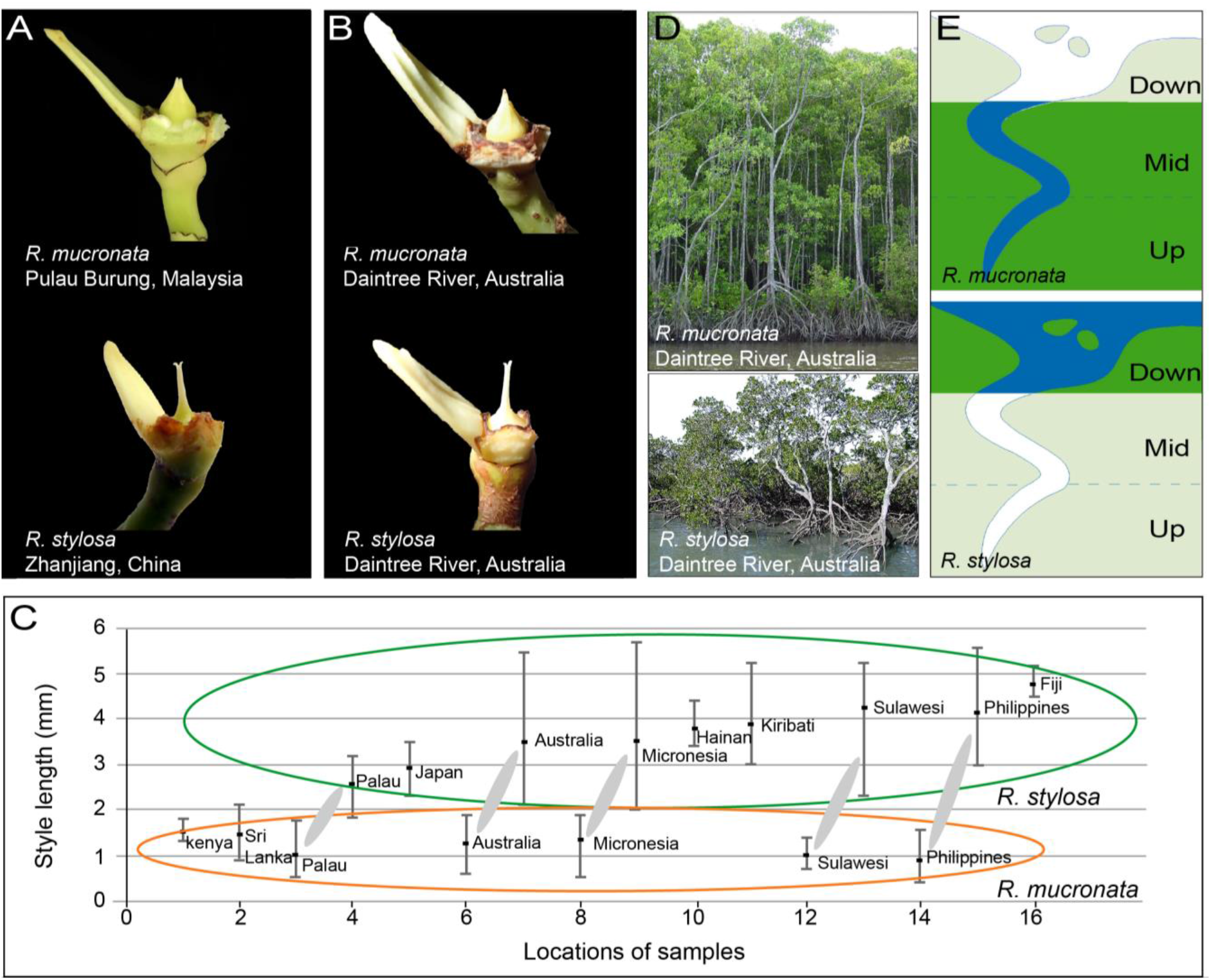
Key diagnostic characters between *R. mucronata* and *R. stylosa*. (A) *R. mucronata* (Pulau Burung, Malaysia, 101°50’14.6’’E, 2°29’33.7’N) and *R. stylosa* (Zhanjiang, China, 109°45’46.50’’E, 21°34’7.32’’N) styles from allopatric sites. (B) *R. mucronata* and *R. stylosa* styles from the Daintree River, Australia, sympatric site. (C) Variation of style lengths of *R. stylosa* (green oval) and *R. mucronata* (orange oval) throughout the Indo-West Pacific region. Solid ellipses contrast sympatric samples. (D) General tree morphology of *R. mucronata* and *R. stylosa* from the Daintree River, Australia, sympatric site. (E) Diagrams showing the habitat preferences of *R. mucronata* and *R. stylosa* in a typical estuary (adapted from mangrove ID (*26*)).

### Characterizations of R. mucronata and R. stylosa in sympatry

We next apply the “secondary sympatry” test to these two species found at the DR site of northern Australia. DR is at the periphery of the distribution of either species (Fig. 2) with *R. mucronata* to the west and *R. stylosa* to the north. It appears that speciation between them had been completed in allopatry and the post-speciation contact happened in DR. Importantly, the two species have remained distinct in their ecology and morphology for a substantial period without intermingling. The two extant species rarely produce F1 hybrids and morphological intermediates are uncommon in our field work. In particular, the style length of each sample is concordant with that of the allopatric populations of the same species (Fig. 3B and Fig. 3C). In the DR area, these two species are parapatric-sympatric with distributions up- or down-river and extensive overlap in the middle (Fig. 3E). This difference in habitat preference is a hallmark of the speciation between *R. mucronata* and *R. stylosa*.

To elaborate on the phylogenetic positions of the DR samples in Fig. 2A, we used the Bayesian clustering analysis implemented in ADMIXTURE (*35*). We identified two genetic components that make up DR sample genomes (Fig. 4A). PCA results also indicate significant admixture in m1 and s1 individuals (Supplementary Fig. S7). Furthermore, because species divergence is monophyletic in all allopatric comparisons, incomplete lineage sorting is an unlikely cause of the observed admixture in the DR samples. In short, we interpret the high *F*_*ST*_ sites as representing divergence after speciation (Fig. 2C) with subsequent admixture in the DR area. Additional tests of introgression (LD analysis, *D* statistic, and the modified *f*_*d*_ statistic) are presented in the Supplement (Tables S8, S10 and Figs. S8-S9).

**Figure 4.**
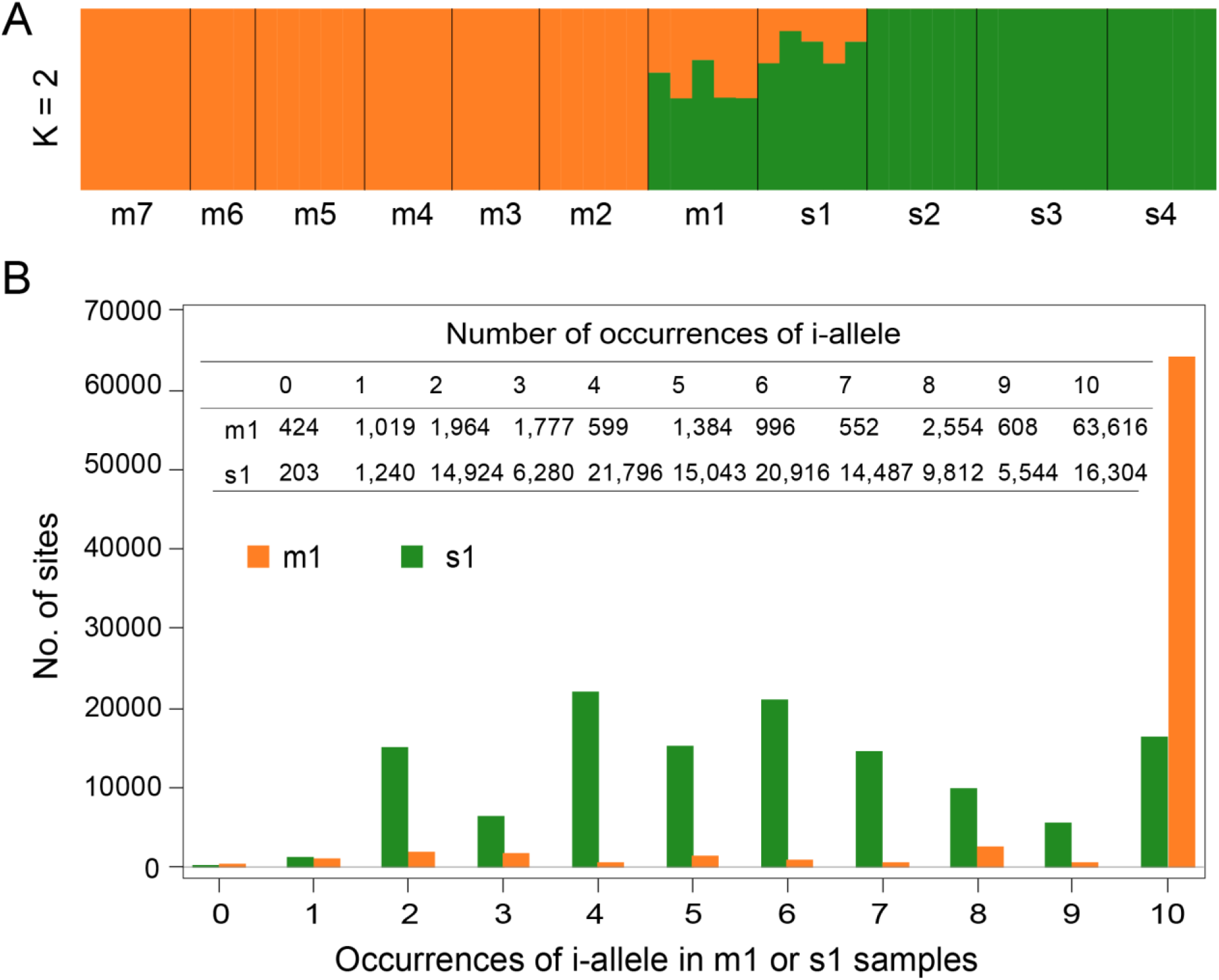
Admixture of genomes between *R. mucronata* and *R. stylosa*. (A) Genetic clustering of all 52 individuals of the two species by ADMIXTURE (K = 2). Orange and green colors denote the *R. mucronata* (m) and *R. stylosa* (s) components, respectively. Individuals from each population are grouped between adjacent black lines. (B) Distribution of i-allele occurrences in m1 (orange) and s1 (green) populations. Given the five individuals (or 10 haploid genomes) from each population, the occurrence ranges from 0 to 10. Exact numbers are given in the inset table (see also Supplement Fig. S10).

### Extensive introgression in sympatry

Using the sympatric samples, we ask the following questions: 1) How many introgressed segments can be found in each species? 2) Is the introgression symmetric? 3) What is the introgressed segment size distribution? A few large blocks are expected if hybridization was recent but many fine-grained blocks should result from old introgressions that have been eroded by recombination. 4) How many genomic segments fail to introgress and what is their genic content? Question 4 will be the subject of the next section.

To identify DR area introgressions, we first define divergent sites (or d-sites) between *R. stylosa* and *R. mucronata* in allopatry. Among the d-sites, we can then define introgressed sites (i-sites) between the m1 and s1 samples in sympatry. There are 305,418 d-sites, defined as sites with *F*_*ST*_ > 0.8 between the two species in allopatry. Note that the bulk of d-sites (212,626) are fully divergent with *F*_*ST*_ = 1.0 (Fig. 2C). An i-site is a locus where the introgressed allele (or i-allele) is found in >= n of the 10 genomes in m1 or s1 sample sets, where n is usually equal to 8. (Note that both m1 and s1 samples have five diploid individuals, or 10 genomes.) In general, if introgression is observed in one direction, say from *R. stylosa* to *R. mucronata* in the DR area, the same site usually does not show introgression (n <= 1) in the reciprocal direction from *R. mucronata* to *R. stylosa*. (In this context, a non-introgressable site, or j-site, is defined as the d-site which contains zero or only one i-allele in both m1 and s1 samples; see Materials and Methods. Note that a small fraction of d-sites are statistically undefinable between i- and j-site.)

To call i-sites, we first define n (the number of genomes carrying the introgressed allele). It is obviously better to set n close to the maximum of 10 for strongly penetrant introgressions. Fig. 4B shows the number of introgressions in the two directions. We set n = 8 for the m1 samples where the i-allele is usually found >= 8 times (orange bars in Fig. 4B). Hence, the results with n= 2 and n = 8 would not be very different. Furthermore, to avoid the confounding presence of remnant ancient polymorphisms, we require introgressions at an i-site to be strongly asymmetric: >= n one way (say, from *R. stylosa* to *R. mucronata*) and <= 1 in the reciprocal direction (Supplementary Fig. S10). I-allele counts are close to uniformly distributed between 2 and 10 (green bars in Fig. 4B) among *R. stylosa* (s1) samples. The asymmetry is probably due to the geography of the DR area, which is at the fringe of the *R. mucronata* distribution. Consequently, gene flow from *R. mucronata* into *R. stylosa* may be more limited here, resulting in the lower frequency of introgressions in the s1 samples. In this regard, setting n = 8 would miss many introgressions in *R. stylosa* leading to a much lower introgression rate than in *R. mucronata*. Nevertheless, the final estimates appear robust even when n is set as low as 2 (see below). Simulations of these scenarios are presented in the Supplement.

Clearly, introgressions do not happen site-by-site, but appear as long segments of DNA consisting of consecutive i-sites. We call these segments “introgression blocks” (or i-blocks) (Supplementary Fig. S11). Fig. 5A shows a segment of the genome that comprises a string of d-sites and i-sites as defined above. These d-sites and i-sites are embedded in a background of low-*F*_*ST*_ or invariant sites shown as dots. This figure shows three i-blocks, each consisting of one, two, or three i-sites. The length of each block is defined by the distance between the two breakpoints flanking it. Unless otherwise specified, we remove the singleton i-blocks that harbor only a single i-site when presenting i-block length distributions.

**Figure 5.**
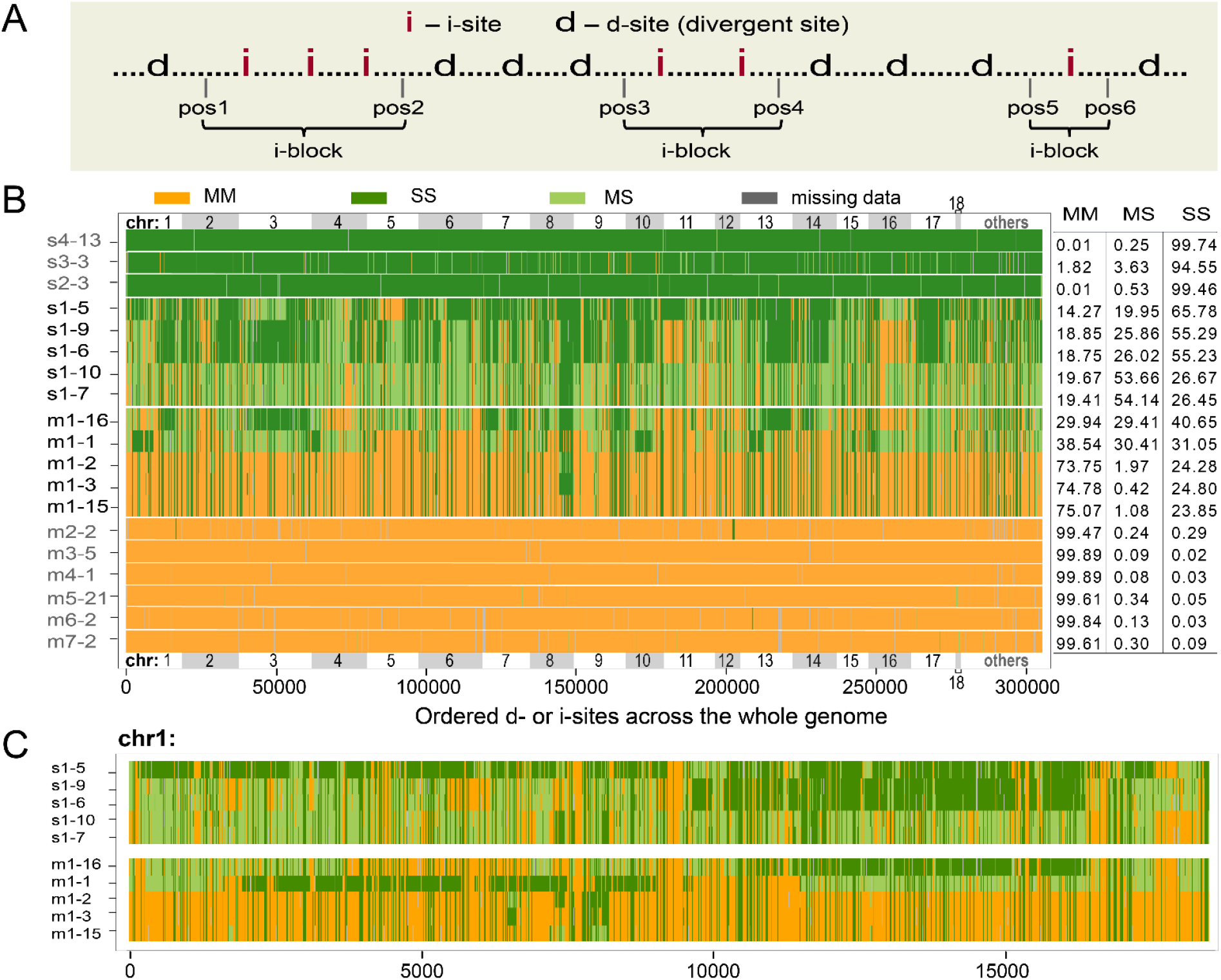
Interspersion between introgressions and non-introgressions. (A) A schematic diagram for delineating i-blocks, each harboring consecutive i-sites without being interrupted by d-sites. The length of an i-block is determined by the midpoints of the flanking (d, i) intervals. Three i-blocks are shown. (B) The genome-wide landscape of i-blocks. Top 18 longest scaffolds (chr1-18) and the rest of the genome (others) are marked and sibling scaffolds are distinguished by gray rectangles. Each row indicates an individual with each vertical line indicating a site. All 10 individuals from the sympatric s1 and m1 populations are shown. For comparison, one individual is randomly selected from each of other populations (see Fig. S12 for the full display). Each site is color-coded for its genotype: MM (orange), MS (light green) and SS (green) type, where M stands for *R. mucronata* and S for *R. stylosa*. The percentages are summarized on the right. Note that extensive interspersions are observed only in the sympatric samples. (C) A close-up view of i-blocks in the longest scaffold (chr1). Only 10 individuals from the sympatric s1 and m1 populations are shown.

The analysis of i-blocks is summarized in Table 1 (see also Supplementary Tables S11-S13). We focus on the results with n = 8 but the results with n = 2 and n = 10 are given for comparison. In the DR area, *R. mucronata* (m1) samples harbor far more introgressions than *R. stylosa* (s1). The bottom of Table 1 at n=8 shows that 16.09 or 23.09% of the *R. mucronata* genomes are introgressions from *R. stylosa*, the two values depending on whether singleton i-blocks are counted. In the opposite direction, 7.97-12.06% of the *R. stylosa* genomes are introgressions. The introgressions in Table 1 are visualized in Figs. 5-6. The salient observation is the highly fine-grained nature of the introgressions. In *R. mucronata*, the introgressions are distributed over 9,963 i-blocks with an average length of 3.24 Kb. In *R. stylosa*, there are 3,874 i-blocks with an average size of 4.13 Kb. Thus, there likely were numerous recombination events that broke introgressions into thousands of tiny i-blocks. It should be noted that Table 1 and Figs. 5-6 present only the extreme cases of introgressions that rise to very high frequencies.

**Table 1.** Summary of high-penetrance introgressed i-blocks between sympatric species

|  | >=2 occurrences of i-allele |  | >=8 occurrences of i-allele |  | =10 occurrences of i-allele |  |
| --- | --- | --- | --- | --- | --- | --- |
|  | m1 | s1 | m1 | s1 | m1 | s1 |
| **No. of i-blocks** | 9,742 | 8,228 | 9,963 | 3,874 | 9,741 | 2,219 |
| **(No. scaffolds with i-blocks)** | (18) | (18) | (18) | (18) | (18) | (18) |
| **Length of i-block (mean)** | 5 bp-944.95 Kb<br>(3.14 Kb) | 8 bp-938.88 Kb<br>(3.49 Kb) | 5 bp-944.95 Kb<br>(3.24 Kb) | 8 bp-938.88 Kb<br>(4.13 Kb) | 5 bp-944.95 Kb<br>(3.02 Kb) | 8 bp-137.48 Kb<br>(4.15 Kb) |
| **No. of i-sites in a block (total number)** | 2-356 bp<br>(40,780) | 2-116 bp<br>(42,862) | 2-349 bp<br>(42,572) | 2-116 bp<br>(19,958) | 2-132 bp<br>(40,727) | 2-100 bp<br>(14,868) |
| **Total length of i-blocks (% of the genome)** | 30.63 Mb<br>(15.26%) | 28.73Mb<br>(14.31%) | 32.29 Mb<br>(16.09%) | 16.00Mb<br>(7.97%) | 29.54 Mb<br>(14.72%) | 9.22Mb<br>(4.59%) |
| **Total length of i-blocks<sup>a</sup> (% of the genome)** | 45.33 Mb<br>(22.58%) | 39.61 Mb<br>(19.74%) | 46.33 Mb<br>(23.09%) | 24.21 Mb<br>(12.06%) | 43.14 Mb<br>(21.49%) | 15.57 Mb<br>(7.76%) |

Note that the species origin of introgressed alleles (i-alleles) is first defined in the allopatric populations. Hence, i-alleles in the DR area can be identified even when they are bi-directional. All i-alleles in this table are uni-directional with, for example, >= 8 in one direction, while the reciprocal direction has <= 1 i-allele. An introgressed block (i-block), unless explicitly stated, has >= 2 introgressed sites (i-sites).
The 18 scaffolds collectively account for 84.38% (200.70 Mb) of the whole genome (237.85 Mb).
<sup>a</sup>These include singleton i-blocks.

The distributions of i-blocks are shown at the large genomic scale in Fig. 5B, at the scaffold scale in Fig. 5C and as individual sites in Fig. 6A-C. Note that only d-sites and i-sites are displayed in these figures. As shown in Fig. 2C, the d- and i-sites are the 305,418 sites with *F*_*ST*_ > 0.8. The rest are invariant or close to invariant sites. The i-blocks are dispersed across the whole genome (Fig. 5B and Supplementary Fig. S12). Indeed, all top 18 scaffolds harbor numerous transitions between i- and d-blocks both in m1 and s1 genomes (Fig. 5B and Table 1). Figure 5C shows that transitions between i- and d-blocks can occur in a few to tens of Kb. At the site level, i-blocks and d-blocks can switch within a small distance (Fig. 6A-6C). An i-block (or d-block) may harbor only one i-site (or d-site), referred to as a singleton block (Fig. 6A-6C, Table 1 and Supplementary Table S11). Singleton blocks, not uncommon but less reliable, are not used in the tally.

**Figure 6.**
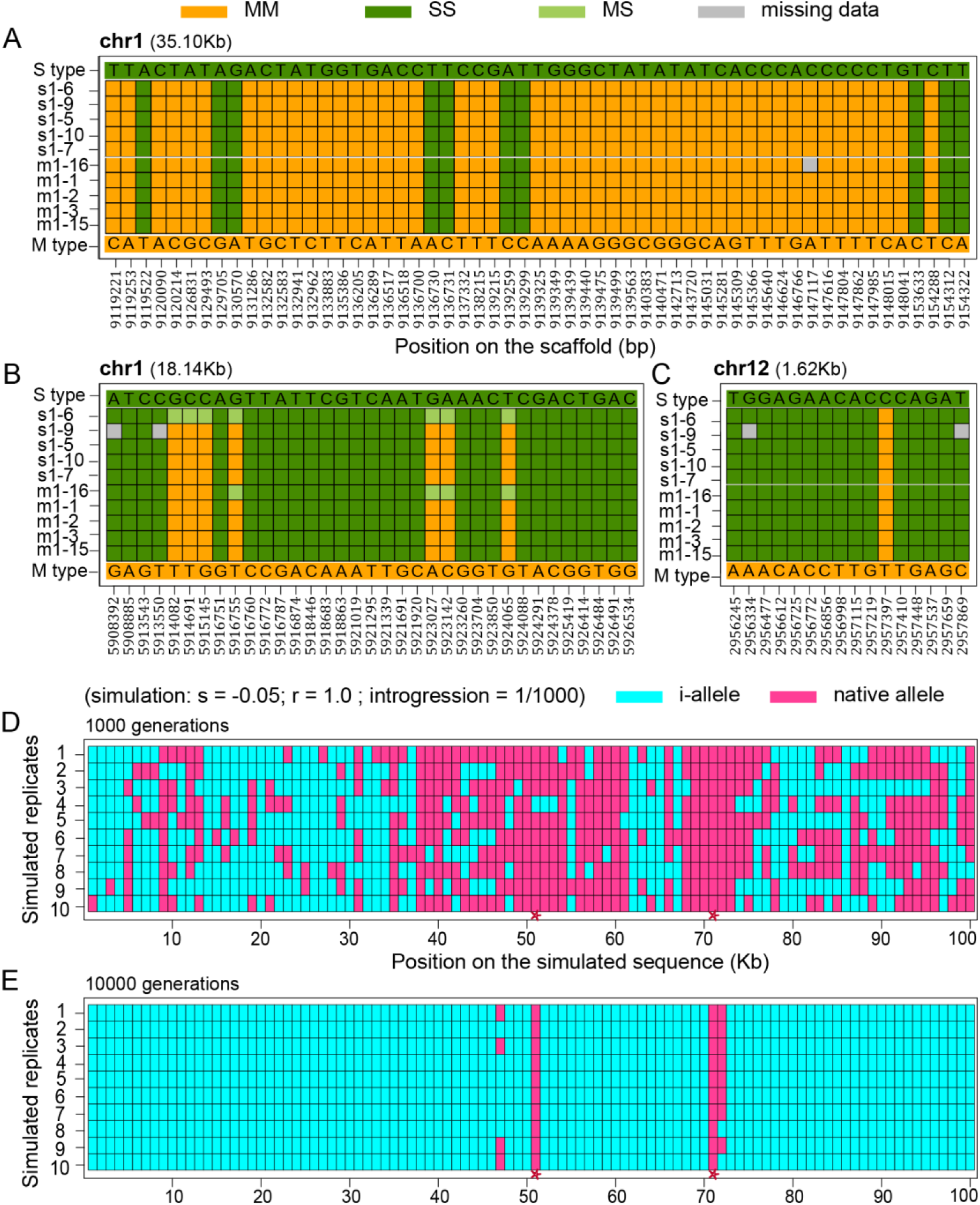
Examples of i-blocks and simulated introgressions in haploid 100Kb genomes. (A-C) Examples of i-blocks in m1 and s1 samples at the site level that show the fine-scale delineation. Color codes are the same as in Fig. 4. (D-E) Simulated introgressions in haploid 100 Kb genomes. This example is from a simulation with strong selection (s = −0.05), high recombination (r = 1.0 for per 100Kb per generation), and low introgression (1/1000 per generation). Two time points are given (see Materials and Methods and Supplementary Fig. S13 for details). Two speciation genes (or loci) under selection at 51 and 71 Kb are marked by red stars at the bottom. Introgressed and non-introgressed sites are marked in blue and pink, respectively. Note that very fine delineations of blocks are possible under the simulated conditions.

The extensive fine-grained introgressions convey two messages. First, hybridizations may happen continually over a long span of time. Each hybridization event would initially bring in whole-chromosome introgressions that are subsequently broken down by recombination. Small DNA fragments may have been introgressed in this piece-meal manner continually. Second, loci underlying differential adaptation between species may be very common such that introgressions tend to be small, and thus free of the introgressed alleles that are deleterious in the genetic background of another species (*36*). In the next section, we will direct our attention toward non-introgressions, which are blocks of native alleles flanked by introgressed DNA segments.

### Very fine-grained interspersion between “introgressable” and “non-introgressable” blocks

Some DNA segments may not be introgressable due to the presence adaptively significant genes. Such loci, by definition, contribute to reproductive isolation or ecological speciation (*5, 37*) and have sometimes been referred to “speciation genes” (*11, 38*–*43*). The number, size, and direction of introgressions therefore depend on several parameters: 1) hybridization rate; 2) the strength of selection against the speciation genes when introgressed; 3) the number and location of speciation loci; 4) recombination rate; and 5) the length of time since initial hybridization.

In a companion study (*6*), we carry out computer simulations based on the Recurrent Selection and Backcross (RSB) model (*44*) (see also Materials and Methods). The RSB model has been proposed for identifying genes underlying complex traits (*44*). It involves repeated dilution the genome of breed A (say, the bulldog) with that of breed B (e.g., the border collie) while retaining the desired phenotypic traits of the former. This is done by continually selecting for the traits of breed A while backcrossing to breed B. The scheme is almost identical to the process of “speciation with gene flow.” They differ only in the parameter values; for example, the length of time in speciation is far greater and gene flow is much smaller, and often bidirectional. The differences necessitate separate simulations for speciation with gene flow. As shown in Fig. 2 of Ref. (*6*) and Fig. 6D-6E in this study (see also Supplementary Fig. S13 and Materials and Methods), introgressions are fine-grained around almost all speciation genes. These patterns resemble the observations reported in this study.

With the j-site defined above, a j-block (i.e., non-introgressable block) is defined as a DNA segment containing at least one j-site (Table 2). Using these stringent criteria, we see ∼1200 j-blocks which together account for < 1% of the genome and harbor 328 coding genes of which 171 contain j-sites (Table 2, Supplementary Table S14 and Fig. S14). For higher confidence, we also show j-blocks with at least two j-sites (Table 2). While only 19 genes containing j-sites are found in these j-blocks (Supplementary Table S15), it is remarkable that six of the 19 genes function in flower development and/or gamete production and development as shown in Table 2 (see the WEGO gene ontology in Supplementary Fig. S14, where a larger set of genes is presented under less stringent criteria). One (*RM_77078*.*7*) of the six genes, known as *EMF1* (embryonic flower 1), regulates reproductive development and is involved in controlling flowering development (*45, 46*). *RM_76773*.*10, RM_76979*.*9* (*NAC2*) and *RM_77530*.*24* are involved in regulating the stamen development, pollen germination and tube growth (*47*–*49*). *RM_76929*.*10, RM_76979* and *RM_77333*.*68* all play a role in embryonic development (*50*–*52*). Since all six genes contain highly differentiated amino acids and non-introgressable sites (j-sites) (Table 2, Supplementary Table S15 and Fig. S15), their involvement in speciation between *R. mucronata* and *R. stylosa* seems plausible.

**Table 2.** High-confidence non-introgressable j-blocks for the identification of genes involved in speciation

|  | >=1 j-sites | >=2 j-sites |
| --- | --- | --- |
| No. of j-blocks (No. scaffolds with j-blocks) | 1,189 (184) | 168 (44) |
| Length of j-blocks - Range (mean) | 3 bp-43.51 Kb (1,010 bp) | 23 bp-35.75 Kb (1,062 bp) |
| No. of j-sites in a block – Range (total j-sites) | 1-6 bp (1,443 bp) | 2-6 bp (422 bp) |
| Total length of j-blocks (% of the genome) | 1,201,823 bp (0.51%) | 178,520 bp (0.075%) |
| No. of genes with j-sites | 171 | 19 |
| No. of genes of flower development with j-sites | -- | 6 (see below) |

| Gene name | j-sites | L(aa) | Site <sup>a</sup> | Function in *Arabidopsis thaliana* |
| --- | --- | --- | --- | --- |
| **RM_76773.10**<br>(AT2G14110) | 3 | 255 | 4 | Haloacid dehalogenase-like hydrolase (HAD) superfamily protein. Participating in pollen germination and tube growth (47). |
| **RM_76929.10**<br>(AT1G55490) | 2 | 294 | 3 | CPN60B. Mutants in this gene develops lesions on its leaves and accelerated cell death to heat shock stress. Participating in embryo and seed development (50). |
| **RM_76979.9**<br>(AT3G15510) | 2 | 199 | 1 | NAC2 (NAC domain containing protein 2). Involved in the regulation of stamen development and embryonic development (48, 51). |
| **RM_77078.7**<br>(AT5G11530) | 3 | 1,415 | 1 | EMF1 (embryonic flower 1). Involved in regulating reproductive development (45, 46). |
| **RM_77333.68**<br>(AT1G08520) | 2 | 755 | 1 | ALB1. Encoding the CHLD subunit of the Mg-chelatase enzyme which involved in chlorophyll biosynthesis. Participating in embryo and seed development (52). |
| **RM_77530.24**<br>(AT3G05420) | 2 | 671 | 2 | ACBP4 (acyl-CoA binding protein 4). Expressed and function in floral lipid metabolism. Playing combinatory roles in pollen development (49). |

A j-block, unless explicitly stated, has >= 2 non-introgressable sites (j-sites).
j-sites: the number of non-introgressable sites within the gene.
L(aa): amino acid sequence length of the gene.
<sup>a</sup> Site: No. of highly differentiated amino acids between *R. mucronata* and *R. stylosa* are given (see Supplementary Fig. S15).

## Discussion

*R. mucronata* and *R. stylosa* in the DR area affirm their species status as they retain their biological characteristics in sympatry. However, we found numerous introgressions where the two species coexist. Since these exchanged segments are small on average, we infer that post-speciation gene flow may have lasted for a long time. There appear to be few exchanges at present with rare F1 hybrids found in DR, as well as at other sampling sites. For example, the m2/s2 collections from Singapore show the expected phylogenetic relationship of their species designation (Fig. 2A and Fig. 2B). Low hybridizations at present have also been reported in Brandan, Indonesia (*27*), Panay Island, Philippines, Kosrae, Micronesia, Yap, Micronesia, and North Sulawesi, Indonesia (*28*).

*R. mucronata* and *R. stylosa* appear to have come into sympatry in the DR area at the right time for extensive post-speciation gene flow to occur. Northern Australia has been suggested as the place where the two species first came into contact (*23*). In this interpretation, the two diverging taxa moved eastward either along the northern coasts of Indian Ocean to Southeast Asia or by crossing the Indian Ocean to Australia (*23*). *R. mucronata* in Southeast Asia then dispersed south and eastward to Australia, while *R. stylosa* in Australia migrated further east and north into SE Asia (see the Supplement) (*23*).

The fine-grained introgressions we find in *R. mucronata* and *R. stylosa* suggest that several conditions during these species’ evolution. First, the diverging populations must have been at the right evolutionary stage when they first came into contact. Had the secondary contact happened earlier, the process of speciation could have been arrested or reversed. Conversely, if the contact would have happened too late, there would be too little gene flow to achieve extensive introgression. The second condition may be even more difficult to satisfy: the two species had to remain in secondary contact for a long time (*4*). This is because numerous recombination events, accumulated over a long period, are necessary to achieve the fine-grained introgression. The third condition is ecological. Sympatric species without niche separation would face competitive exclusion (*53*). *R. mucronata* and *R. stylosa* had evolved a degree of niche separation that results in incomplete overlap in habitat preference (Fig. 3E).

The fine-grained pattern of introgression between *R. mucronata* and *R. stylosa* we find is exceptional in the literature. Nevertheless, the rarity of such observations does not necessarily mean that post-speciation gene flow is unusual. It only means that the opportunities for such observations may be rare. Indeed, *R. mucronata* and *R. stylosa* in the DR area represent the confluence of the three conditions presented above. Post-speciation gene flow could not be easily detected without careful planning.

In this study, we interpret BSC to require complete RI as proposed in the original literature (*3*) (see Wu (*5*) for further analyses). Although many have argued that BSC should tolerate “a little” gene flow (textbooks like Futuyma’s (*1*)), the argument is conceptually inconsistent with BSC. It also makes RI an operationally undefinable quantity. After all, genomic studies have suggested that more than a third of the genome may be exchangeable between species (*5, 6, 11, 17, 54*). In conclusion, BSC and the full RI should be abandoned as a key criterion for species delineation given the many recent genomic studies (*11, 21, 36, 55*–*59*) and the observations reported here.

## Acknowledgements

We thank Loren Rieseberg, James Mallet, Matthew Hahn, Richard Abbott, Patrik Nosil, Jeffrey Feder, Trevor Price, Nicholas Barton and Nolan Kane for insightful comments. We thank Wei Lun Ng for the photo of *R. mucronata* style in Fig. 2A. This study was supported by the National Natural Science Foundation of China (91731301, 31600182 and 31830005); the National Key Research and Development Plan (2017FY100705); Guangdong Basic and Applied Basic Research Foundation (2019A1515010752); the China Postdoctoral Science Foundation (2019M663212) and the Chang Hungta Science Foundation of Sun Yat-Sen University.

## Data Availability

Genomic raw reads of *R. mucronata* and *R. stylosa* individuals have been deposited to the Genome Sequence Archive of National Genomics Data Center &BIG Data Center with GSA accession number CRA001688 (http://bigd.big.ac.cn/gsa).

## Materials and Methods

### Plant material, genome sequencing and assembly

*R. mucronata* and *R. stylosa* individuals were collected for whole-genome sequencing from Dongzhai Harbor, Hainan, China (110°35’5.79’’ E, 19°56’39.67’’ N, Supplementary Table S2), although the *R. mucronata* individual was originally introduced from Australia (*60*). Genomic DNA extraction from leaves was done following the CTAB method (*61*). Total RNA was extracted from leaves, flowers, and fruits using the modified CTAB method (*62*). Short-read libraries were constructed and sequenced using the BGISEQ-500 platform. 50 Kb long-read libraries were prepared using the 10X Genomics (Illumina Hiseq X Ten) platform. Before assembling, low-quality reads, adaptor sequences, N and polyA contamination were filtered out. The 10X Genomics data was used to assemble a draft genome using SUPERNOVA (*63*). To calibrate and refine the assembly, we used clean short reads and Hi-C reads based on the 3D-DNA (3D DNA de novo genome assembly) pipeline (https://github.com/theaidenlab/3d-dna). After manual check and calibration with Juicebox (https://github.com/aidenlab/Juicebox), we anchored 18 pseudo-chromosomes (chr1-18) both for *R. mucronata* (200.70 Mb, or 84.38%) and *R. stylosa* (219.66 Mb, or 86.63%) genomes, corresponding to their diploid chromosome number of 2n = 36 (Supplementary Table S1) (*64, 65*). The transcriptome data was used to perform protein-coding gene annotation (26,540 genes in *R. mucronata* and 30,375 in *R. stylosa*; Supplementary Table S1). Genome completeness (94.2% for *R. mucronta* and 94.3% for *R. stylosa*) was assessed using the lineage database eudicotyledons_odb10 and the BUSCO software (https://github.com/c-omics/busco, Supplementary Table S1). Raw reads were mapped to the de novo genomes using the Burrows-Wheeler Aligner (BWA) (*66, 67*). Mapping rates were as high as 94.52% for *R. mucronta* and 92.74% for *R. stylosa* (Supplementary Table S2).

### Collinearity analysis

We performed collinearity analyses within and between *R. mucronata* and *R. stylosa* genomes. Alignment was performed on protein sequences using BLASTP (with a cutoff e-value of 10^−10^, identity ≥ 40%). We then used MCScanX (*68*) to identify syntenic or collinear blocks, with blocks containing at least five paired homologous genes accepted. Genomic distribution of collinear blocks was visualized using the Circos (v0.65) software (Fig. 1B and Supplementary Figs. S1-S2) (*69*). 267 and 305 syntenic blocks were identified within the *R. mucronata* and *R. stylosa* genome, occupying up to 32.01% (8,496 of all 26,540 genes) and 29.61% (8,995 of all 30,375 genes) genes of each genome (Supplementary Figs. S1-S2). Through collinearity analysis between *R. mucronata* and *R. stylosa*, we found 663 inter-specific collinear blocks with 18,705 *R. mucronata* (70.48% of 26,540 genes) and 18,952 *R. stylosa* genes (62.39% of 30,375 genes) involved (Fig. 1B).

### Gene family analysis and divergence time estimation

To estimate divergence times, we used the de novo genomes of *Carallia longipes* (unpublished data), *Bruguiera gymnorrhiza* (*29*), *Rhizophora apiculate* (*22*), *R. mucronata*, and *R. stylosa*. We used OrthoFinder-2.2.7 (*70*) to identify gene families (*67, 71*). All proteins from these five species were merged to perform an all-to-all alignment using DIAMOND (with a cutoff e-value of 10^−30^) (*72*). 20,971 gene families were identified among the five species, 6,014 containing only one gene in each species (i.e., single-copy orthologs). Genes from the five species that fell into shared single-copy orthologs were aligned using MUSCLE (*73*) and codon sequences were obtained using PAL2NAL.v14 (*74*). We used JMODELTEST2 (*75*) to select an appropriate nucleotide-substitution model for reconstructing the phylogeny. To reconstruct the phylogenetic tree, we used RAXML (*30*) and IQTREE (*31*) with the best-fit model (GTR+G) and 1000 bootstrap replicates (Supplementary Fig. S3). Finally, the program MCMCTREE from the PAML4.8 package (*76*) was employed to estimate divergence times (Fig. 1C, Supplementary Table S3 and Fig. S3), using “seq like (usedata = 1)”, “JC69 (model = 0, alpha = 0)” and “independent rates (clock = 2)”. The time constraints were based on the earliest known fossil records of mangrove lineages (hypocotyl fossils of *Bruguiera* in early Eocene (constraint 1: 47.8-56 Myr ago) and the oldest records of *Rhizophora* in the upper Eocene (constraint 2:33.9-38.0 Myr ago) (*22, 77*–*79*).

### Sampling and genome re-sequencing

To make the samples of *R. mucronata* and *R. stylosa* more representative, we collected individuals both in allopatry and sympatry in the Indo-West Pacific region (Fig. 2 and Supplementary Tables S4). We re-sequenced 31 *R. mucronata* individuals from seven populations and 21 *R. stylosa* individuals from four populations (Fig. 2 and Supplementary Tables S4-S6). To tell apart the two species by morphology, we observed the style length and shape in the bud and took photos (Fig. 3 and Supplementary Tables S9). Fresh leaves were sampled from individual trees and dried with silica gel. Genomic DNA extraction was done following the CTAB method (*61*). Short-read libraries were sequenced using the Illumina Hiseq 2000 platform with insert size of 350bp and constructed following the TruSeq DNA Sample Preparation Guide. We obtained high quality sequence data for each individual genome, with coverage in the 12 to 22X range (Supplementary Tables S5-S6).

### SNP calling and genetic diversity detection

To identify variants, clean reads from all 52 individuals were mapped to the de novo *R. mucronata* genome using the Burrows-Wheeler Aligner (BWA) (*66*). SAMtools (*80*) were used to import, sort, and pair bam files and remove duplications. To obtain high quality variants, single nucleotide polymorphisms (SNPs) were called and filtered using the Genome Analysis Toolkit (GATK) (*81*) and the SAMtools/bcftools (*80*) pipeline. Only consensus SNPs called by both pipelines were retained for downstream analyses. To remove low-quality variants, we eliminated all loci that had base quality (Q) or mapping quality (q) smaller than 20. We additionally performed the following stringent filtering: 1) at least two reads had to support the minor allele to call a heterozygote; 2) only homozygous SNPs with read depth >=2 were retained. After filtering, we selected these high-quality sites for further analyses, with multi-allelic (>=3) sites, insertions, and deletions excluded. To estimate genetic diversity in each population, we calculated θ_w_ (Watterson’s θ_w_) and θ_π_ (Nei and Li’s θ_π_) (*82, 83*) within each population (Supplementary Table S4). To estimate genomic divergence between *R. mucronata* and *R. stylosa* populations, we calculated the genetic differentiation coefficient (*F*_*ST*_) (Fig. 2D) (*82, 84*).

### Detecting gene flow

We applied Patterson’s *D* statistic and a modified *f*_*d*_ statistic to quantify gene flow (*85, 86*). A positive *D* or *f*_*d*_ value is an indicator of introgression (Supplement Fig. S8 and Table S10). The basic model has three ingroups (P_1_, P_2_, and P_3_) and the outgroup (O) in the genealogical relationship (((P_1_, P_2_), P_3_), O). In our analysis, P_1_ and P_2_ are different populations from the same species *R. mucronata* (or *R. stylosa*), while P_3_ corresponds to the other species. The outgroup is *R. apiculate* (*22*). Positive *D* values imply that P_2_ and P_3_ have more shared alleles than P_1_ and P_3_ (see Supplement Table S10 and Fig. S9). The plink-1.07 (*87*) software package was used to estimate linkage disequilibrium (LD), represented by the r^2^ statistic within each population or group (Supplement Fig. S8). LD decay was used to test for the presence of admixture events. We also calculated LD decay in sympatric populations in Singapore (s2 and m2) and allopatric *R. mucronata* and *R. stylosa* populations as controls (Supplement Fig. S8).

### Genomic scan for introgressed and non-introgressable blocks

We used four predefined taxa: m1 (*R. mucronata* population in Daintree River), s1 (*R. stylosa* population in Daintree River), M_allo_ (allopatric *R. mucronata* populations m2-m7), and S_allo_ (allopatric *R. stylosa* populations s2-s4). To get a more informative data set, we filtered sites with too many missing genotypes in each taxon or low divergence (*F*_*ST*_ <= 0.8) between M_allo_ and S_allo_. We retained 305,418 SNPs (*F*_*ST*_ > 0.8) which we call divergent sites or d-sites between M_allo_ and S_allo_ (Fig. 2C). 212,626 of the d-sites are fixed (*F*_*ST*_ = 1.0 and *D*_*xy*_ = 1.0) between M_allo_ and S_allo_ (Fig. 2C). There are four possible states of each d-site: homozygous *R. mucronata* variant (MM), homozygous *R. stylosa* variant (SS), heterozygote (MS), or missing data.

We then looked for introgressed sites (i-sites) and non-introgressable sites (j-sites) among all the d-sites across m1 and s1 genomes. We have five diploid individuals (or 10 haploid genomes) from the m1 and s1 populations. We have defined allele classes as follows. **Introgressed allele** (**i-allele)**: *R. stylosa* variant in m1 populations or *R. mucronata* variant in s1 populations. **i-site:** an i-site in m1 or in s1 genomes is defined as >= 8 occurrences of i-allele out of the 10 genomes (Fig. 4B and Supplementary Fig. S10). **j-site:** a d-site with <=1 occurrences of i-allele in both m1 and s1 populations (Fig. 4B and Supplementary Fig. S10). **i-block:** A genomic block in one species is considered introgressed from the other species if one or more i-sites continuously (without disruption by other d-sites) are present (Fig. 5A, Fig. 6A-C and Supplementary Fig. S11). The length of an i-block is determined by the midpoint between the flanking (d-sites, i-sites) intervals (as shown in Fig. 5A). **j-block:** a genomic block with one or more j-sites continuously. We define the boundaries the same as for i-blocks.

### Simulations of genomic sequences under hybridization, selection, and recombination

To probe the influences of hybridization, selection, and recombination on genomic sequences, we carried out computer simulations based on the Recurrent Selection and Backcross (RSB) model (*44*). We set high and low levels for each parameter. Population size was set at 1000. The length of simulated sequences was 100 Kb (for convenience, 1 Kb is the basic unit that cannot be separated by recombination). The original allele in the sequence and an i-allele from the other species were differentially labeled. Hence, at the beginning of the simulations, the sequences of all individuals were in original allelic states (100 x). After several generations of hybridization, selection, and recombination, the sequences become shuffled (Fig. 6D-E and Fig. S13).

We first set a low hybridization rate (introgression or migration rate, m = 0.001 per generation) and recombination (10E-6 per generation between adjacent base pairs). For every generation, 999 individuals were picked from the original population and one from the other population (or species). The recombination probability (r) for a 100 Kb sequence was about 0.1. Since population size is 1000, there will be an average of 100 individuals with recombination in each generation. Two loci under negative selection (#51 and #71) were defined in the simulated sequences. If one or both loci harbor an i-allele, the relative fitness of this sequence is 0.95 (Fig. S13A) or 0.99 (Fig. S13B). We also examined a high introgression rate regime (10/1000). In this case, four loci (#41, #51, #71 and #76) were negatively selected (relative fitness = 0.95 for an i-allele) (Fig. S13C). The scenarios in Fig. S13A show that a lower recombination rate (r = 0.1) increases the size of non-introgressed DNA segments, because the neutral genes near positions 51 and 71 were selected against along with the speciation loci. Figure S13B-C shows that a reduced selection intensity (s = −0.01) or a 10-fold higher introgression rate give rise to extensive introgressions. Interestingly, partial introgressions were detected even at positions 51 and 71, where selection acts against the invading alleles.

Finally, we simulated genomic sequences under a high recombination rate (10E-5, r = 1.0 for a 100 Kb simulated sequence per generation) and a low introgression rate (1/1000 per generation). Two loci (#51 and #71) were negatively selected (relative fitness = 0.95 for an i-allele) (Fig. 6D-E and Supplementary Fig. S14D-F). The simulations suggest that, given the right parameter values (Fig. 6D-E and Supplementary Fig. S14D-F), the pattern of introgression would follow exactly the prediction based solely on selection, whereby only the alleles of the speciation loci cannot be introgressed. The rest of the genome, even right next to the speciation loci, is freely shared between species.

## References

1. D. J. Futuyma, M. Kirkpatrick, Evolution (Sunderland, MA: Sinauer Associates, Inc., 2017).

2. T. Dobzhansky, Genetics and the Origin of Species (Columbia University Press, New York, 1937).

3. E. Mayr, Animal Species and Evolution (Cambridge, MA: Harvard University Press, 1963).

4. J. A. Coyne, H. A. Orr, Speciation (Sunderland, MA: Sinauer Associates, Inc., 2004).

5. C.-I Wu, The genic view of the process of speciation. J. Evol. Biol. 14, 851–865 (2001).

6. X. Wang, Z. He, S. Shi, C.-I Wu, Genes and speciation: Is it time to abandon the biological species concept? Natl. Sci. Rev. 7, 1387–1397 (2020).

7. C.-I Wu, A. W. Davis, Evolution of Postmating Reproductive Isolation: The Composite Nature of Haldane’s Rule and Its Genetic Bases. Am. Nat. 142, 187–212 (1993).

8. K. Sawamura, A. W. Davis, C.-I Wu, Genetic analysis of speciation by means of introgression into Drosophila melanogaster. Proc. Natl. Acad. Sci. U. S. A. 97, 2652–2655 (2000).

9. M. F. Palopoli, C.-I Wu, Genetics of hybrid male sterility between Drosophila sibling species: A complex web of epistasis is revealed in interspecific studies. Genetics. 138, 329–341 (1994).

10. E. L. Cabot, A. W. Davis, N. A. Johnson, C.-I Wu, Genetics of reproductive isolation in the Drosophila simulans clade: Complex epistasis underlying hybrid male sterility. Genetics. 137, 175–189 (1994).

11. C.-I Wu, C. T. Ting, Genes and speciation. Nat. Rev. Genet. 5, 114–122 (2004).

12. J. I. Meier, D. A. Marques, S. Mwaiko, C. E. Wagner, L. Excoffier, O. Seehausen, Ancient hybridization fuels rapid cichlid fish adaptive radiations. Nat. Commun. 8, 14363 (2017).

13. O. Seehausen, Patterns in fish radiation are compatible with Pleistocene desiccation of Lake Victoria and 14 600 year history for its cichlid species flock. Proc. R. Soc. B Biol. Sci. 269, 491–497 (2002).

14. M. J. Ryan, Food, song and speciation. Nature. 409, 139–140 (2001).

15. S. Lamichhaney et al., Evolution of Darwin’s finches and their beaks revealed by genome sequencing. Nature. 518, 371–375 (2015).

16. T. L. Turner, M. W. Hahn, S. V. Nuzhdin, Genomic islands of speciation in Anopheles gambiae. PLoS Biol. 3, 1572–1578 (2005).

17. S. H. Martin et al., Genome-wide evidence for speciation with gene flow in Heliconius butterflies. Genome Res. 23, 1817–1828 (2013).

18. R. Abbott et al., Hybridization and speciation. J. Evol. Biol. 26, 229–246 (2013).

19. J. W. Poelstra et al., The genomic landscape underlying phenotypic integrity in the face of gene flow in crows. Science 344, 1410–1414 (2014).

20. N. Vijay et al., Evolution of heterogeneous genome differentiation across multiple contact zones in a crow species complex. Nat. Commun. 7, 13195 (2016).

21. Z. He et al., Speciation with gene flow via cycles of isolation and migration: Insights from multiple mangrove taxa. Natl. Sci. Rev. 6, 275–288 (2019).

22. S. Xu et al., The origin, diversification and adaptation of a major mangrove clade (Rhizophoreae) revealed by whole-genome sequencing. Natl. Sci. Rev. 4, 721–734 (2017).

23. N. C. Duke, E. Lo, M. Sun, Global distribution and genetic discontinuities of mangroves - Emerging patterns in the evolution of Rhizophora. Trees - Struct. Funct. 16, 65–79 (2002).

24. N. C. Duke, in Elevitch CR (ed) Traditional trees of Pacific Islands: their culture, environment, and use (Permanent Agriculture Resources, Holualoa, Hawaii, 2006), Pp.641–660.

25. N. C. Duke, M. C. Ball, J. C. Ellison, Factors influencing biodiversity and distributional gradients in mangroves. Glob. Ecol. Biogeogr. Lett. 7, 27–47 (1998).

26. N. C. Duke, “World Mangrove iD: expert information at your fingertips” App Store Version 1.2,July 2017. MangroveWatch Publication, Australia - e-book. (2017).

27. A. K. S. Wee et al., Genetic differentiation and phylogeography of partially sympatric species complex Rhizophora mucronata Lam. and R. stylosa Griff. using SSR markers Phylogenetics and phylogeography. BMC Evol. Biol. 15, 1–13 (2015).

28. Y.-B. Yan, N. C. Duke, M. Sun, Comparative Analysis of the Pattern of Population Genetic Diversity in Three Indo-West Pacific Rhizophora Mangrove Species. Front. Plant Sci. 7, 1–17 (2016).

29. Z. He et al., Convergent adaptation of the genomes of woody plants at the land-sea interface. Natl. Sci. Rev. 7, 978–993 (2020).

30. A. Stamatakis, RAxML version 8: A tool for phylogenetic analysis and post-analysis of large phylogenies. Bioinformatics. 30, 1312–1313 (2014).

31. L. T. Nguyen, H. A. Schmidt, A. Von Haeseler, B. Q. Minh, IQ-TREE: A fast and effective stochastic algorithm for estimating maximum-likelihood phylogenies. Mol. Biol. Evol. 32, 268–274 (2015).

32. S. Kumar, G. Stecher, K. Tamura, MEGA7: Molecular Evolutionary Genetics Analysis Version 7.0 for Bigger Datasets. Mol. Biol. Evol. 33, 1870–4 (2016).

33. K. J. Galinsky, G. Bhatia, P. R. Loh, S. Georgiev, S. Mukherjee, N. J. Patterson, A. L. Price, Fast Principal-Component Analysis Reveals Convergent Evolution of ADH1B in Europe and East Asia. Am. J. Hum. Genet. 98, 456–472 (2016).

34. D. J. Futuyma, Evolution (Sunderland, MA: Sinauer Associates, Inc., 2005).

35. D. H. Alexander, J. Novembre, K. Lange, Fast model-based estimation of ancestry in unrelated individuals. Genome Res. 19, 1655–64 (2009).

36. S. Fang et al., Incompatibility and competitive exclusion of genomic segments between sibling Drosophila species. PLoS Genet. 8, e1002795 (2012).

37. D. Schluter, Evidence for ecological speciation and its alternative. Science 323, 737–741 (2009).

38. D. C. Presgraves, The molecular evolutionary basis of species formation. Nat. Rev. Genet. 11, 175–180 (2010).

39. J. Mallet, What does Drosophila genetics tell us about speciation? Trends Ecol. Evol. 21, 386–393 (2006).

40. S. Sun, C. T. Ting, C.-I Wu, The normal function of a speciation gene, Odysseus, and its hybrid sterility effect. Science 305, 81–83 (2004).

41. C. T. Ting, S. C. Tsaur, M. L. Wu, C.-I Wu, A rapidly evolving homeobox at the site of a hybrid sterility gene. Science 282, 1501–1504 (1998).

42. P. Nosil, D. Schluter, The genes underlying the process of speciation. Trends Ecol. Evol. 26, 160–167 (2011).

43. J. Wittbrodt et al., Novel putative receptor tyrosine kinase encoded by the melanoma-inducing Tu locus in Xiphophorus. Nature. 341, 415–421 (1989).

44. Z. W. Luo, C.-I Wu, M. J. Kearsey, Precision and high-resolution mapping of quantitative trait loci by use of recurrent selection, backcross or intercross schemes. Genetics. 161, 915–929 (2002).

45. R. Sánchez, M. Y. Kim, M. Calonje, Y. H. Moon, Z. R. Sung, Temporal and spatial requirement of EMF1 activity for arabidopsis vegetative and reproductive development. Mol. Plant. 2, 643–653 (2009).

46. H. Y. Park, S. Y. Lee, H. Y. Seok, S. H. Kim, Z. R. Sung, Y. H. Moon, EMF1 interacts with EIP1, EIP6 or EIP9 involved in the regulation of flowering time in Arabidopsis. Plant Cell Physiol. 52, 1376–1388 (2011).

47. Y. Wang, W.-Z. Zhang, L.-F. Song, J.-J. Zou, Z. Su, W.-H. Wu, Transcriptome Analyses Show Changes in Gene Expression to Accompany Pollen Germination and Tube Growth in Arabidopsis. Plant Physiol. 148, 1201–1211 (2008).

48. A. Mandaokar et al., Transcriptional regulators of stamen development in Arabidopsis identified by transcriptional profiling. Plant J. 46, 984–1008 (2006).

49. A. S. Hsiao, E. C. Yeung, Z. W. Ye, M. L. Chye, The Arabidopsis cytosolic Acyl-CoA-binding proteins play combinatory roles in pollen development. Plant Cell Physiol. 56, 322–333 (2015).

50. X. Ke et al., Functional divergence of chloroplast Cpn60α subunits during Arabidopsis embryo development. PLoS Genet. 13, e1007036 (2017).

51. T. Kunieda et al., NAC family proteins NARS1/NAC2 and NARS2/NAM in the outer integument regulate embryogenesis in arabidopsis. Plant Cell. 20, 2631–2642 (2008).

52. N. Bryant, J. Lloyd, C. Sweeney, F. Myouga, D. Meinke, Identification of Nuclear Genes Encoding Chloroplast-Localized Proteins Required for Embryo Development in Arabidopsis. Plant Physiol. 155, 1678–1689 (2010).

53. G. Hardin, The competitive exclusion principle. Science 131, 1292–1297 (1960).

54. H. Ellegren et al., The genomic landscape of species divergence in Ficedula flycatchers. Nature. 491, 756–760 (2012).

55. O. Seehausen, Hybridization and adaptive radiation. Trends Ecol. Evol. 19, 198–207 (2004).

56. L. H. Rieseberg, A new model of speciation. Natl. Sci. Rev. 6, 289–290 (2019).

57. R. J. Abbott, A mixing-isolation-mixing model of speciation can potentially explain hotspots of species diversity. Natl. Sci. Rev. 6, 290–291 (2019).

58. T. Price, Allo-parapatric speciation goes offshore. Natl. Sci. Rev. 6, 289–289 (2019).

59. N. H. Barton, Is speciation driven by cycles of mixing and isolation? Natl. Sci. Rev. 6, 291–292 (2019).

60. B. Liao, S. Zheng, Y. Chen, M. Li, W. Zeng, D. Zheng, Preliminary Report on Introduction of Several Alien Mangrove Plants in Dongzhai Harbour of Hainan Provbince. J. Cent. SOUTH For. Univ. 26, 63–67 (2006).

61. J. J. Doyle, A rapid DNA isolation procedure for small quantities of fresh leaf tissue. Phytochem. Bull. 19, 11–15 (1987).

62. G. Yang, R. Zhou, T. Tang, S. Shi, Simple and efficient isolation of high-quality total RNA from Hibiscus tiliaceus, a mangrove associate and its relatives. Prep. Biochem. Biotechnol. 38, 257–264 (2008).

63. N. I. Weisenfeld, V. Kumar, P. Shah, D. M. Church, D. B. Jaffe, Direct determination of diploid genome sequences. Genome Res. 27, 757–767 (2017).

64. A. P. Tyagi, Cytogenetics and reproductive biology of mangroves in Rhizophoraceae. Aust. J. Bot. 50, 601–605 (2002).

65. D. Subramanian, Cytological Studies of some Mangroove Flora of Tamilnadu. Cytologia (Tokyo). 53, 87–92 (1988).

66. H. Li, Aligning sequence reads, clone sequences and assembly contigs with BWA-MEM. http://arxiv.org/abs/1303.3997 (2013).

67. G. D. Wang et al., Structural variation during dog domestication: Insights from gray wolf and dhole genomes. Natl. Sci. Rev. 6, 110–122 (2019).

68. Y. Wang et al., MCScanX: A toolkit for detection and evolutionary analysis of gene synteny and collinearity. Nucleic Acids Res. 40, e49 (2012).

69. M. Krzywinski et al., Circos: An information aesthetic for comparative genomics. Genome Res. 19, 1639–1645 (2009).

70. D. M. Emms, S. Kelly, OrthoFinder: solving fundamental biases in whole genome comparisons dramatically improves orthogroup inference accuracy. Genome Biol. 16, 157 (2015).

71. T. Lin et al., Genome analysis of Taraxacum kok-saghyz Rodin provides new insights into rubber biosynthesis. Natl. Sci. Rev. 5, 78–87 (2018).

72. B. Buchfink, C. Xie, D. H. Huson, Fast and sensitive protein alignment using DIAMOND. Nat. Methods. 12, 59–60 (2015).

73. R. C. Edgar, MUSCLE: A multiple sequence alignment method with reduced time and space complexity. BMC Bioinformatics. 5, 113 (2004).

74. M. Suyama, D. Torrents, P. Bork, PAL2NAL: Robust conversion of protein sequence alignments into the corresponding codon alignments. Nucleic Acids Res. 34, W609–W612 (2006).

75. D. Darriba, G. L. Taboada, R. Doallo, D. Posada, JModelTest 2: More models, new heuristics and parallel computing. Nat. Methods. 9, 772 (2012).

76. Z. Yang, PAML 4: Phylogenetic analysis by maximum likelihood. Mol. Biol. Evol. 24, 1586–1591 (2007).

77. A. Graham, Paleobotanical Evidence and Molecular Data in Reconstructing the Historical Phytogeography of Rhizophoraceae 1. Ann. Missouri Bot. Gard. 93, 325–334 (2006).

78. M. E. Collinson, Fossil Plants of the London Clay (Palaeontological Association Field Guide to Fossils, 1983).

79. J. Muller, Fossil pollen records of extant angiosperms. Bot. Rev. 47, 1–142 (1981).

80. H. Li et al., The Sequence Alignment/Map format and SAMtools. Bioinformatics. 25, 2078–2079 (2009).

81. A. McKenna et al., The Genome Analysis Toolkit: a MapReduce framework for analyzing next-generation DNA sequencing data. Genome Res. 20, 1297–303 (2010).

82. M. Nei, Analysis of gene diversity in subdivided populations. Proc. Natl. Acad. Sci. U. S. A. 70, 3321–3323 (1973).

83. M. Nei, W. H. Li, Mathematical model for studying genetic variation in terms of restriction endonucleases. Proc. Natl. Acad. Sci. U. S. A. 76, 5269–5273 (1979).

84. S. Wright, The genetic structure of populations. Ann. Eugenetics. 16, 97–159 (1951).

85. E. Y. Durand, N. Patterson, D. Reich, M. Slatkin, Testing for ancient admixture between closely related populations. Mol. Biol. Evol. 28, 2239–2252 (2011).

86. R. E. Green et al., A draft sequence of the neandertal genome. Science 328, 710–722 (2010).

87. S. Purcell et al., PLINK: a tool set for whole-genome association and population-based linkage analyses. Am. J. Hum. Genet. 81, 559–75 (2007).

