## Supplementary materials for "In the absence of reproductive isolation – Extensive gene flow after speciation"

### Supplementary information for
### In the absence of reproductive isolation – Extensive gene flow after speciation

Xinfeng Wang [a,*], Zixiao Guo [a,*], Ziwen He [a,*], Shaohua Xu [a], Shao Shao [a], Sen Li [a], Ming Yang [a], Qipian Chen [a], Cairong Zhong [b], Zhongyi Wu [a], Norman C. Duke [c], Suhua Shi [a†]

[a] State Key Laboratory of Biocontrol, Guangdong Key Lab of Plant Resources, School of Life Sciences, Sun Yat-Sen University, Guangzhou, Guangdong, China

[b] Hainan Dongzhai Harbor National Nature Reserve Administration, Haikou, Hainan, China

[c] Centre for Tropical Water and Aquatic Ecosystem Research, James Cook University, Townsville, Queensland, Australia

* These authors contributed equally to this work

**Supplementary information**

**This file includes:**

Supplementary Notes

Tables S1-S15

Figures S1-S15

References

**Supplementary Text**

In this study, the de novo sequenced *Rhizophora mucronata* and *R. stylosa* individual are abbreviated as RM and RS (Table S2), respectively. *R. mucronata* populations are labeled as m1, m2, m3, m4, m5, m6, and m7, while *R. stylosa* populations as s1, s2, s3, and s4, where "m" stands for *R. mucronata* and “s” stands for *R. stylosa* (Table S4). To refer to all the *R. stylosa* or *R. mucronata* populations, we use "$S_{all}$" or "$M_{all}$". We use "$S_{allo}$" to refer to the allopatric *R. stylosa* populations s2-s4 and "$M_{allo}$" for the allopatric *R. mucronata* populations m2-m7.

***Additional introgression tests between sympatric species***

i) When comparing the two species, we identified 325 fixed and 900,474 shared SNPs (Table S8). Removing m1 and s1 (i.e., the DR samples), changes the SNP counts to 215,350 and 259,759 respectively. Removing allopatric populations does not affect the locus counts much, and the numbers remain around 350 and 860,000 (Table S8). This suggests that incomplete lineage sorting as not the cause of the observed admixture in the DR samples.

ii) We also observed an increased genome-wide linkage disequilibrium (LD) in m1 and s1 (Fig. S8). Hence, if the background admixture in the allopatric populations reflects ancient polymorphism (or incomplete lineage sorting), the excess admixture we observed in m1 and s1 can be reasonably attributed to introgression private to m1 and s1.

iii) We further used the ABBA-BABA (or *D*) statistic to test for the excess of shared derived alleles due to introgression (*85*, *86*). A positive “*D*” or “$f_d$” value is an indicator of introgression (gene flow). The tests with m1 and s1 as the subject branches all showed significant positive “*D*” and “$f_d$” values ($P<0.01$, Table S10). In contrast, no mean *D* values significantly deviated from 0 ($P>0.5$) in tests without m1 or s1 (Table S10). Using 500kb sliding windows, we constructed distributions of *D* values across the genome. Between 97.80% and 99.76% of the windows had positive *D* values in the tests with m1 and s1 (Fig. S9). In other words, introgression was only found between m1 and s1 locally in Daintree River, where *R. stylosa* and *R. mucronata* are found together.

***Detecting highly differentiated amino acids between R. mucronata and R. stylosa***

To find highly differentiated amino acids in the 19 candidate genes in j-blocks, we obtained protein sequences of all 52 individuals for each gene (Table 2 and Table S15). We used the following criteria to call highly differentiated amino acids: 1) the differentiated nucleotide in the codon is a non-synonymous site and with $F_{ST} > 0.8$ between $M_{allo}$ and $S_{allo}$ samples; 2) there are no identical homozygotes between *R. mucronata* and *R. stylosa*. We found 32 such amino acids between *R. mucronata* and *R. stylosa* in the 19 genes (Table S15). Twelve of these sites are in the six genes involved in flower development and/or gamete production and development (Table 2 and Fig. S15).

***The possible evolutionary trajectory and speciation of R. mucronata and R. stylosa***

These observations re-enforce the notion that species characters vary over geographic range. Accepting that, we then ask what additional circumstances would be needed for speciation to occur and how might these conditions appear for the two *Rhizophora* species in this case study?

The characters shown for the two closely related species by these genetic and morphological studies have revealed distinct patterns and traits that appear to follow an ordered series of features tending towards genetic isolation and speciation. The order of events may have followed a scenario similar to the following (*23*):

1) It started with the dispersal of Asian *R. mucronata* into a vacant habitat as conditions became suitable in the west and south, but with concurrent selection favoring drift towards arid and marine traits to differentiate East African *R. mucronata.*
2) Propagules were transported due to their exceptional ability for long distance dispersal and a specialized capability for establishment and growth in marine coastal conditions in a broad range of wet and dry climatic conditions.
3) These circumstances would have transported populations south and east towards Australia (in one or more founder events that selected a subset of genotypes which at some point became *R. stylosa*).
4) The longer style of *R. stylosa* further implies some possibly greater reliance on a particular pollinator, but this has not been established.
5) Ancestral Australian *R. stylosa* migrates further east and north along northern Australian shorelines and then into SE Asia as well as the western Pacific where at some point it re-unites with ancestral Asian *R. mucronata* (still ecologically conditioned for upstream brackish estuarine locations).
6) Before this event, Asian *R. mucronata* populations would have been expanding east and south as conditions became more suitable across the region, but the proximity and size of populations would likely have prevented significant isolation events and no further speciation would have occurred.
7) The geomorphic circumstances of continental drift and their timing were likely critical in the progress of the biological events of dispersal and speciation (*88*).

#### Supplementary Tables:

##### Table S1 *De novo* sequencing and assembly of *R. mucronata* and *R. stylosa* genomes

| | ***R. mucronata*** | ***R. stylosa*** |
|---|---|---|
| Estimated genome size (Mb) [a] | 306.91 | 365.44 |
| Total assembly size (Mb) | 237.85 | 253.54 |
| Total number of contigs | 54,203 | 80,721 |
| Total size (Mb) | 248.51 | 263.44 |
| The longest contig (Mb) | 3.14 | 3.85 |
| No. contigs (length >=1 Mb) | 37 | 16 |
| Contig N50 length (Kb) | 344.17 | 105.85 |
| Total number of scaffolds | 14,496 | 9,750 |
| The longest scaffold (Mb) | 16.96 | 19.13 |
| The shortest scaffold (bp) | 1,000 | 1,000 |
| No. scaffolds (length >= 5 Mb) | 18 | 19 |
| Total length of the top 18 longest scaffolds (Mb) | 200.70 | 219.66 |
| Proportion of the assembly genome [b] | 84.38% | 86.63% |
| Scaffold N50 length (Mb) | 12.03 | 12.59 |
| Scaffold N50 count | 9 | 9 |
| Total N (Mb) | 14.72 | 24.56 |
| GC content (%) | 35.46 | 36.56 |
| No. annotation genes | 26,540 | 30,375 |
| Repeat content (%) | 30.22 | 20.67 |
| Genome completeness [c] | 94.2% (1,999 of 2,121 groups) | 94.3% (2,000 of 2,121 groups) |
| Protein completeness [c] | 90.9% (1,927 of 2,121 groups) | 92.4% (1,959 of 2,121 groups) |

[a] Estimated genome size by flow cytometry/K-mer analysis.

[b] Proportion occupied by the top 18 longest scaffolds of the assembly genome.

c Genomic/protein completeness was assessed with the lineage database eudicotyledons_odb10, using BUSCO (https://github.com/c-omics/busco).

#### Table S2 *De novo* sequencing information and genetic statistics of the *R. mucronata* and *R. stylosa* genomes

| | ***R. mucronata*** | ***R. stylosa*** |
|---|---|---|
| Sampling location (Longitude, latitude) | Dongzhai Harbor, Hainan, China (110°35'5.79" E, 19°56'39.67" N) | Dongzhai Harbor,Hainan, China (110°35'5.79" E, 19°56'39.67" N) |
| Sample ID | RM | RS |
| Read length (bp) | 100 | 100 |
| Raw read pairs | 169,210,696 | 180,644,954 |
| Retained clean reads pairs | 141,318,480 | 153,186,318 |
| Mapped reads | 267,136,374 | 284,127,598 |
| Reads mapping rate (%) | 94.52 | 92.74 |
| Assembly genome size (bp) | 237,845,386 | 253,544,496 |
| Effective sites [a] | 196,713,446 | 198,044,661 |
| Coverage rate [b] | 0.83 | 0.78 |
| Mean depth [c] | 142 | 142 |
| No. of heterozygotes | 94,170 | 106,490 |
| Ht [d] (per Kb) | 0.48 | 0. 56 |
| No. private variants | 37,923 | 48,766 |
| No. shared variants | 46,218 | |
| **$D_{xy}$ (RM vs. RS)** | **0.0031** | |

[a] Effective sites: the sites that have at least two reads mapped (depth>=2) for each de novo sequencing individual.

[b] Coverage rate: the proportion occupied by effective sites of the assembly genome size.

[c] Mean depth is (raw reads*reads length)/assembly genome size.

[d] Mean genome-wide heterozygosity (Ht): the proportion of all effective sites occupied by heterozygotes.

#### Table S3 Divergence time and credible interval of each node of the Rhizophoreae group using *MCMCTREE*

| Node | Divergence time (Mya) | 95%HPD [a] (Mya) |
|---|---|---|
| 1 | 3.75 | [3.15, 4.35] |
| 2 | 10.28 | [8.93, 11.65] |
| 3 | 40.15 | [37.51, 43.50] |
| 4 | 57.03 | [52.24, 60.56] |

[a] 95%HPD: 95% highest posterior density credible interval of each node is shown. Nodes are marked in Supplementary Fig. S3b. Mya: million years ago.

#### Table S4 Sampling, re-sequencing information, and genetic diversity statistics for *R. mucronata* and *R. stylosa* populations

| Sampling location | Longitude, latitude | Pop ID | Sample size | Effective sites [a] | Coverage rate [b] | SNPs (x$10^5$) | $\theta_w$ (Kb) | $\theta_\pi$ (Kb) |
|---|---|---|---|---|---|---|---|---|
| ***R. mucronata*** | | | | | | | | |
| Daintree River, Australia | 145°26'16.28" E,16°17'12.44" S | m1 | 5 | 1.98E+08 | 0.83 | 7.80 | 1.39 | 1.37 |
| Saint John's Island, Singapore | 103°50'30.19" E, 1°13'6.60" N | m2 | 5 | 1.92E+08 | 0.81 | 2.81 | 0.52 | 0.65 |
| Chai-Ya, Thailand | 99°15'32.35" E, 9°21'23.07" N | m3 | 4 | 1.91E+08 | 0.80 | 2.02 | 0.41 | 0.49 |
| Ranong, Thailand | 98°37'26.01" E, 9°57'36.26" N | m4 | 4 | 1.95E+08 | 0.82 | 5.12 | 1.01 | 1.17 |
| Tanjung Piai, Malaysia | 103°21'1.86" E, 1°24'8.11" N | m5 | 5 | 1.95E+08 | 0.82 | 3.53 | 0.64 | 0.74 |
| Mauritius | 57°41'18.33" E, 20°20'26.37" S | m6 | 3 | 1.90E+08 | 0.80 | 0.86 | 0.20 | 0.24 |
| Kenya | 39°36'1.82" E, 4°24'24.15" S | m7 | 5 | 1.98E+08 | 0.83 | 1.90 | 0.34 | 0.42 |
| ***R. stylosa*** | | | | | | | | |
| Daintree River, Australia | 145°26'16.28" E, 16°17'12.44" S | s1 | 5 | 1.95E+08 | 0.82 | 9.05 | 1.64 | 2.09 |
| Saint John's Island, Singapore | 103°50'30.19" E, 1°13'6.60" N | s2 | 5 | 1.90E+08 | 0.80 | 1.28 | 0.24 | 0.30 |
| Darwin, Australia | 130°54'22.64" E, 12°25'5.75" S | s3 | 6 | 1.89E+08 | 0.80 | 5.99 | 1.05 | 1.23 |
| Hainan, China | 110°35'5.79" E, 19°56'39.67" N | s4 | 5 | 1.93E+08 | 0.81 | 1.19 | 0.22 | 0.28 |

[a] Effective sites: the sites that have at least one individual mapped (depth>=2) for each population.

[b] Coverage rate: the proportion occupied by effective sites of the reference genome size. The reference (*R. mucronata*) genome size is 237,845,386 bp.

#### Table S5 Re-sequencing characteristics and heterozygosity of each *R. mucronata* genome

| Population ID | Ind. ID | Read length (bp) | Raw read pairs (E+07) | Clean read pairs (E+07) | Mapped reads (E+07) | Reads mapping rate (%) | Effective sites [a] (E+08) | Coverage rate [b] | Mean Depth [c] | Ht [d] (per Kb) |
|---|---|---|---|---|---|---|---|---|---|---|
| **m1** (Daintree River, Australia) | m1-1 | 125 | 2.03 | 2.03 | 3.58 | 88.33 | 1.93 | 0.81 | 19X | 2.00 |
| | m1-15 | 125 | 1.37 | 1.37 | 2.51 | 91.32 | 1.89 | 0.79 | 13X | 0.47 |
| | m1-16 | 125 | 1.36 | 1.36 | 2.40 | 88.09 | 1.90 | 0.80 | 13X | 2.09 |
| | m1-2 | 100 | 2.18 | 1.99 | 3.68 | 92.45 | 1.78 | 0.75 | 15X | 0.50 |
| | m1-3 | 100 | 2.51 | 2.28 | 4.21 | 92.28 | 1.81 | 0.76 | 18X | 0.38 |
| **m2** (Saint John's Island, Singapore) | m2-1 | 150 | 1.49 | 1.43 | 2.56 | 89.71 | 1.82 | 0.76 | 16X | 0.80 |
| | m2-2 | 150 | 2.09 | 1.98 | 3.53 | 89.02 | 1.85 | 0.78 | 22X | 0.42 |
| | m2-3 | 150 | 1.62 | 1.58 | 2.96 | 93.90 | 1.81 | 0.76 | 19X | 0.60 |
| | m2-4 | 150 | 1.44 | 1.40 | 2.64 | 94.42 | 1.83 | 0.77 | 17X | 0.65 |
| | m2-5 | 150 | 1.78 | 1.73 | 3.25 | 93.79 | 1.85 | 0.78 | 20X | 0.66 |
| **m3** (Chai-Ya, Thailand) | m3-12 | 125 | 1.48 | 1.48 | 2.75 | 92.55 | 1.84 | 0.77 | 14X | 0.46 |
| | m3-14 | 125 | 1.62 | 1.62 | 3.03 | 93.75 | 1.83 | 0.77 | 16X | 0.50 |
| | m3-5 | 125 | 1.51 | 1.51 | 2.85 | 94.07 | 1.86 | 0.78 | 15X | 0.46 |
| | m3-7 | 125 | 1.42 | 1.42 | 2.65 | 93.38 | 1.84 | 0.77 | 14X | 0.50 |
| **m4** (Ranong, Thailand) | m4-1 | 125 | 1.70 | 1.70 | 3.16 | 92.91 | 1.85 | 0.78 | 17X | 0.45 |
| | m4-2 | 125 | 1.46 | 1.46 | 2.71 | 92.79 | 1.85 | 0.78 | 14X | 0.63 |
| | m4-3 | 125 | 1.65 | 1.65 | 3.08 | 93.13 | 1.83 | 0.77 | 16X | 0.62 |
| | m4-4 | 125 | 1.49 | 1.49 | 2.76 | 92.86 | 1.89 | 0.79 | 15X | 0.67 |
| **m5** (Tanjung Piai, Malaysia) | m5-12 | 125 | 1.40 | 1.34 | 2.49 | 93.02 | 1.90 | 0.80 | 13X | 0.85 |
| | m5-14 | 125 | 1.90 | 1.90 | 3.33 | 87.41 | 1.82 | 0.76 | 17X | 0.44 |
| | m5-21 | 125 | 2.01 | 2.01 | 3.60 | 89.43 | 1.79 | 0.75 | 19X | 0.65 |
| | m5-32 | 125 | 1.34 | 1.28 | 2.38 | 92.90 | 1.79 | 0.75 | 13X | 0.84 |
| | m5-9 | 125 | 1.27 | 1.19 | 2.20 | 92.88 | 1.78 | 0.75 | 12X | 0.52 |
| **m6** (Mauritius) | m6-1 | 125 | 1.35 | 1.35 | 2.47 | 91.44 | 1.77 | 0.74 | 13X | 0.38 |
| | m6-2 | 125 | 1.34 | 1.34 | 2.51 | 93.88 | 1.83 | 0.77 | 13X | 0.34 |
| | m6-3 | 125 | 1.32 | 1.32 | 2.42 | 91.90 | 1.82 | 0.76 | 13X | 0.35 |
| **m7** (Kenya) | m7-10 | 125 | 1.55 | 1.55 | 2.79 | 89.77 | 1.84 | 0.78 | 15X | 0.51 |
| | m7-11 | 125 | 1.48 | 1.48 | 2.59 | 87.58 | 1.85 | 0.78 | 14X | 0.52 |
| | m7-2 | 125 | 1.77 | 1.77 | 3.11 | 87.57 | 1.83 | 0.77 | 16X | 0.41 |
| | m7-7 | 125 | 2.11 | 2.11 | 3.75 | 88.80 | 1.90 | 0.80 | 20X | 0.52 |
| | m7-9 | 125 | 1.92 | 1.92 | 3.38 | 87.74 | 1.83 | 0.77 | 18X | 0.53 |

[a] Effective sites: all sites that have at least two reads mapped (depth>=2) in each site.

[b] Coverage rate: the proportion occupied by effective sites of the reference genome size. The reference (*R. mucronata*) genome size equals to 237,845,386 bp.

[c] Mean depth is (mapped reads*reads length)/reference genome size.

[d] Mean genome-wide heterozygosity (Ht): the proportion of all effective sites occupied by heterozygotes.

#### Table S6 Re-sequencing characteristics and heterozygosity of each *R. stylosa* genome

| Population ID | Ind. ID | Read length (bp) | Raw read pairs (E+07) | Clean read pairs (E+07) | Mapped reads (E+07) | Reads mapping rate (%) | Effective sites [a] (E+08) | Coverage rate [b] | Mean Depth [c] | Ht [d] (per Kb) |
|---|---|---|---|---|---|---|---|---|---|---|
| **s1** (Daintree River, Australia) | **s1-** | 100 | 1.83 | 1.68 | 3.11 | 92.80 | 1.85 | 0.78 | 13X | 3.32 |
| | **s1-5** | 100 | 1.72 | 1.58 | 2.91 | 92.24 | 1.89 | 0.80 | 12X | 1.58 |
| | **s1-6** | 100 | 1.95 | 1.79 | 3.27 | 91.31 | 1.85 | 0.78 | 14X | 2.03 |
| | **s1-7** | 100 | 2.01 | 1.85 | 3.42 | 92.74 | 1.78 | 0.75 | 14X | 3.43 |
| | **s1-9** | 100 | 1.81 | 1.66 | 3.07 | 92.20 | 1.82 | 0.77 | 13X | 2.04 |
| **s2** (Saint John's Island, Singapore) | **s2-1** | 150 | 1.58 | 1.54 | 2.88 | 93.93 | 1.77 | 0.74 | 18X | 0.40 |
| | **s2-3** | 150 | 1.72 | 1.68 | 3.17 | 94.57 | 1.80 | 0.76 | 20X | 0.40 |
| | **s2-4** | 150 | 1.53 | 1.48 | 2.76 | 93.18 | 1.80 | 0.76 | 17X | 0.39 |
| | **s2-5** | 150 | 1.41 | 1.38 | 2.60 | 94.05 | 1.81 | 0.76 | 16X | 0.38 |
| | **s2-6** | 150 | 1.43 | 1.39 | 2.64 | 94.61 | 1.81 | 0.76 | 17X | 0.40 |
| **s3** (Darwin, Australia) | **s3-1** | 100 | 2.24 | 2.07 | 3.76 | 91.04 | 1.78 | 0.75 | 16X | 1.27 |
| | **s3-2** | 100 | 2.19 | 2.01 | 3.59 | 89.47 | 1.80 | 0.76 | 15X | 1.23 |
| | **s3-3** | 100 | 1.94 | 1.80 | 3.24 | 90.07 | 1.81 | 0.76 | 14X | 0.78 |
| | **s3-4** | 100 | 2.16 | 1.97 | 3.46 | 87.64 | 1.83 | 0.77 | 15X | 1.18 |
| | **s3-5** | 100 | 2.26 | 2.07 | 3.74 | 90.38 | 1.79 | 0.75 | 16X | 1.26 |
| | **s3-6** | 100 | 2.69 | 2.46 | 4.46 | 90.68 | 1.81 | 0.76 | 19X | 1.31 |
| **s4** (Hainan, China) | **s4-** | 100 | 1.99 | 1.82 | 3.33 | 91.42 | 1.80 | 0.76 | 14X | 0.40 |
| | **s4-** | 100 | 2.00 | 1.84 | 3.34 | 90.85 | 1.79 | 0.75 | 14X | 0.37 |
| | **s4-3** | 100 | 1.85 | 1.69 | 3.12 | 91.98 | 1.79 | 0.75 | 13X | 0.37 |
| | **s4-4** | 100 | 1.79 | 1.65 | 3.04 | 92.24 | 1.76 | 0.74 | 13X | 0.39 |
| | **s4-7** | 100 | 1.91 | 1.75 | 3.22 | 91.79 | 1.86 | 0.78 | 14X | 0.35 |

[a]Effective sites: all sites that have at least two reads mapped (depth>=2) in each site.

[b] Coverage rate: the proportion occupied by effective sites of the reference genome size. The reference (*R. mucronata*) genome size equals to 237,845,386 bp.

[c] Mean depth is (mapped reads*reads length)/reference genome size.

[d] Mean genome-wide heterozygosity (Ht): the proportion of all effective sites occupied by heterozygotes.

**Table S7 Genome-wide genetic divergence between and within *R. mucronata* and *R. stylosa* populations**

| Description | Relationship | Mean genome-wide $D_{xy}$ (per Kb) | Mean genome-wide $F_{ST}$ |
|---|---|---|---|
| Sympatric divergence | **m1 vs. s1** | **2.40** | **0.21** |
| Genetic divergence within *R. mucronata* | m1 vs. $M_{allo}$ [a] | 3.41 | 0.26 |
| Genetic divergence within *R. stylosa* | s1 vs. $S_{allo}$ [b] | 2.90 | 0.25 |
| Genetic divergence between interspecific populations | m1 vs. $S_{allo}$ | 3.94 | 0.43 |
| | s1 vs. $M_{allo}$ | 3.90 | 0.27 |
| | **$M_{allo}$ vs. $S_{allo}$** | **4.37** | **0.49** |
| Genetic divergence between *R. mucronata* and *R. stylosa* | **$M_{all}$ vs. $S_{all}$** [c] | **4.14** | **0.34** |

[a] "$M_{allo}$": allopatric *R. mucronata* populations m2-m7.

[b] "$S_{allo}$": allopatric *R. stylosa* populations s2-s4.

[c] "$M_{all}$" represents all *R. mucronata* populations m1-m7, while "$S_{all}$" represents all *R. stylosa* populations s1-s4.

**Table S8 *R. mucronata* and *R. stylosa* polymorphism statistics**

| Removed populations [a] | Fixed difference | Shared polymorphisms | Private polymorphisms in *R. stylosa* | Private polymorphisms in *R. mucronata* | Total SNPs |
|---|---|---|---|---|---|
| none | 325 | 900,474 | 322,965 | 517,259 | 1,741,023 |
| **m1, s1** | **215,350** | **259,759** | **492,260** | **590,503** | **1,741,023** |
| m2, s2 | 343 | 885,312 | 314,701 | 518,424 | 1,741,023 |
| m2, s3 | 418 | 845,759 | 207,212 | 557,977 | 1,741,023 |
| m2, s4 | 341 | 886,514 | 311,177 | 517,222 | 1,741,023 |
| m3, s2 | 342 | 887,920 | 312,093 | 523,907 | 1,741,023 |
| m3, s3 | 417 | 848,259 | 204,712 | 563,568 | 1,741,023 |
| m3, s4 | 340 | 889,211 | 308,480 | 522,616 | 1,741,023 |
| m4, s2 | 370 | 881,242 | 318,771 | 459,051 | 1,741,023 |
| m4, s3 | 445 | 843,004 | 209,967 | 497,289 | 1,741,023 |
| m4, s4 | 366 | 882,372 | 315,319 | 457,921 | 1,741,023 |
| m5, s2 | 351 | 885,605 | 314,408 | 513,202 | 1,741,023 |
| m5, s3 | 422 | 846,239 | 206,732 | 552,568 | 1,741,023 |
| m5, s4 | 346 | 886,887 | 310,804 | 511,920 | 1,741,023 |
| m6, s2 | 346 | 886,636 | 313,377 | 518,285 | 1,741,023 |
| m6, s3 | 420 | 847,372 | 205,599 | 557,549 | 1,741,023 |
| m6, s4 | 343 | 887,946 | 309,745 | 516,975 | 1,741,023 |
| m7, s2 | 356 | 881,564 | 318,449 | 504,829 | 1,741,023 |
| m7, s3 | 432 | 844,254 | 208,717 | 542,139 | 1,741,023 |
| m7, s4 | 354 | 882,873 | 314,818 | 503,520 | 1,741,023 |

[a] Removed populations: we removed two populations from all samples each time and then calculate the polymorphisms in the rest. *R. mucronata* and *R. stylosa*. "none" means we kept all samples.

**Table S9 Additional diagnostic morphological features to identify *R. mucronata* and *R. stylosa***

| **Feature** | ***R. mucronata*** | ***R. stylosa*** |
|---|---|---|
| **bracts and bracteoles** | minute bracts and bracteoles | distinct bracts and bracteoles |
| **inflorescences** | 1-2 flowered inflorescences | 4-16 flowered inflorescences |
| **flower buds** | irregular obovoid closed flower buds | regular ovoid-elliptic closed flower buds |
| **propagules** | long propagules reaching ~80 cm | ~60 cm |

**Table S10 Patterson's *D* statistic and improved $f_d$ statistic, showing evidence of gene flow between *R. mucronata* (m1) and *R. stylosa* (s1) in sympatry in Daintree River, Australia**

| Model code [a] | Pop1 [b] | Pop2 [b] | Pop3 [b] | Outgr-oup [b] | *D* ± std err [c] | Z-score | P-value | $f_d$ ± std err [d] |
|---|---|---|---|---|---|---|---|---|
| **1** | m7 | m6 | s1 | ra | -0.0272 ± 0.0151 | -0.0890 | 0.929 | -0.00517 ± 4.54E-05 |
| **2** | m7 | m6 | s2 | ra | 0.0138 ± 0.0216 | 0.0316 | 0.975 | 0.00284 ± 7.78 E-05 |
| **3** | m7 | m6 | s3 | ra | -0.0136 ± 0.0206 | -0.0327 | 0.974 | -0.00226 ± 7.70E-05 |
| **4** | m7 | m6 | s4 | ra | -0.00350 ± 0.0216 | -0.00798 | 0.994 | 0.00161 ± 7.64E-05 |
| **5** | m3 | m2 | s2 | ra | -0.00127 ± 0.0135 | -0.00466 | 0.996 | -0.00200 ± 2.57E-05 |
| **6** | m4 | m2 | s2 | ra | 0.113 ± 0.0140 | 0.399 | 0.690 | 0.0108 ± 4.85E-05 |
| **7** | m5 | m2 | s2 | ra | 0.00626 ± 0.0120 | 0.0257 | 0.979 | -0.000231 ± 5.06E-06 |
| **8** | m6 | m2 | s2 | ra | -0.169 ± 0.0209 | -0.401 | 0.688 | -0.0333 ± 1.61E-04 |
| **9** | m7 | m2 | s2 | ra | -0.174 ± 0.0209 | -0.412 | 0.681 | -0.0309 ±2.16E-04 |
| **10** | s3 | s2 | m2 | ra | 0.128 ± 0.0180 | 0.351 | 0.726 | 0.0409 ± 1.71E-04 |
| **11** | s4 | s2 | m2 | ra | 0.00597 ± 0.0209 | 0.0141 | 0.989 | -0.0137 ± 1.10E-04 |
| **12** | m2 | m1 | s1 | ra | 0.640 ± 0.00861 | 3.67*** | **2.42E-04** | 0.445 ± 7.23E-05 |
| **13** | m3 | m1 | s1 | ra | 0.637 ± 0.00894 | 3.52*** | **4.32E-04** | 0.442 ± 7.43E-05 |
| **14** | m4 | m1 | s1 | ra | 0.646 ± 0.0101 | 3.17*** | **1.51E-03** | 0.447 ±7.77E-05 |
| **15** | m5 | m1 | s1 | ra | 0.642 ± 0.00832 | 3.81*** | **1.38E-04** | 0.445 ± 7.36E-05 |
| **16** | m6 | m1 | s1 | ra | 0.565 ± 0.0103 | 2.70*** | **6.89E-03** | 0.422 ± 8.39E-05 |
| **17** | m7 | m1 | s1 | ra | 0.561 ± 0.0102 | 2.71*** | **6.65E-03** | 0.420 ± 8.87E-05 |
| **18** | s2 | s1 | m1 | ra | 0.607 ± 0.00949 | 3.16*** | **1.59E-03** | 0.548 ± 1.14E-04 |
| **19** | s3 | s1 | m1 | ra | 0.600 ± 0.00912 | 3.24*** | **1.18E-03** | 0.553 ± 8.49E-05 |
| **20** | s4 | s1 | m1 | ra | 0.605 ± 0.00951 | 3.14*** | **1.67E-03** | 0.544 ± 9.83E-05 |

[a]: code of *D* statistic models. Models 1-11 exclude sympatric populations m1 and s1; models 12-20 contain sympatric populations m1 and s1.

[b]: Pop1, Pop2, Pop3 and Outgroup respectively refer to the three ingroups and the outgroup (ra: *R. apiculata*) following the genealogical relationship (((Pop1, Pop2), Pop3), Outgroup).

[c]: *D* statistic, given as a ratio *D* ± standard error.

***: the genome-wide average *D*-statistic value *D* is significantly derived from 0 with P < 0.01, indicating the existence of gene flow between m1 and s1 populations.

[d]: $f_d$ statistic, given as an admixed proportion $f_d$ ± standard error.

**Table S11 Introgressed site (i-site) distribution across introgressed blocks (or i-blocks) in m1 and s1 genomes**

| **The i-sites range of i-blocks** | **>=2 occurrences of i-allele** | | **>=4 occurrences of i-allele** | | **>=6 occurrences of i-allele** | | **>=8 occurrences of i-allele** | | **=10 occurrences of i-allele** | |
|---|---|---|---|---|---|---|---|---|---|---|
| | **m1** | **s1** | **m1** | **s1** | **m1** | **s1** | **m1** | **s1** | **m1** | **s1** |
| 1 (singleton block) | 17961 | 8912 | 17195 | 8023 | 17246 | 7226 | 17278 | 5411 | 17091 | 3333 |
| 2 | 4937 | 3004 | 4878 | 2790 | 4957 | 2341 | 4959 | 1572 | 4858 | 917 |
| 3 | 1944 | 1563 | 1954 | 1478 | 2001 | 1167 | 2003 | 740 | 1936 | 401 |
| 4 | 945 | 973 | 971 | 922 | 993 | 709 | 985 | 416 | 966 | 241 |
| 5 | 527 | 611 | 535 | 604 | 540 | 451 | 549 | 261 | 544 | 159 |
| 5-10 | 861 | 1286 | 878 | 1323 | 915 | 997 | 907 | 529 | 883 | 292 |
| 10-15 | 241 | 412 | 247 | 399 | 256 | 332 | 260 | 160 | 259 | 97 |
| 15-20 | 109 | 148 | 110 | 140 | 114 | 144 | 113 | 65 | 121 | 28 |
| 20-30 | 93 | 118 | 95 | 135 | 96 | 128 | 93 | 74 | 89 | 44 |
| 30-40 | 33 | 73 | 34 | 63 | 35 | 62 | 36 | 28 | 33 | 21 |
| 40-50 | 20 | 21 | 20 | 21 | 20 | 25 | 18 | 18 | 20 | 10 |
| 50-60 | 8 | 6 | 9 | 7 | 9 | 6 | 11 | 3 | 12 | 4 |
| 60-70 | 6 | 3 | 6 | 5 | 6 | 3 | 8 | 2 | 3 | 3 |
| 70-80 | 4 | 2 | 4 | 2 | 4 | 3 | 4 | 1 | 5 | 0 |
| 80-90 | 2 | 2 | 2 | 1 | 2 | 1 | 2 | 1 | 3 | 0 |
| 90-100 | 4 | 4 | 5 | 4 | 5 | 4 | 4 | 3 | 3 | 2 |
| >100 | 8 | 2 | 10 | 2 | 11 | 3 | 11 | 1 | 6 | 0 |
| Total blocks | 27703 | 17140 | 26953 | 15919 | 27210 | 13602 | 27241 | 9285 | 26832 | 5552 |
| Blocks (>=2 i-sites) | 9742 | 8228 | 9758 | 7896 | 9964 | 6376 | 9963 | 3874 | 9741 | 2219 |

#### Table S12 Length distribution of introgressed blocks (i-blocks) in m1 and s1 genomes

| **Length range of i-blocks** | **>=2 occurrences of i-allele** | | **>=4 occurrences of i-allele** | | **>=6 occurrences of i-allele** | | **>=8 occurrences of i-allele** | | **=10 occurrences of i-allele** | |
|---|---|---|---|---|---|---|---|---|---|---|
| | **m1** | **s1** | **m1** | **s1** | **m1** | | **m1** | **s1** | **m1** | **s1** |
| 1-10bp | 217 | 96 | 219 | 80 | 221 | 62 | 219 | 43 | 214 | 28 |
| 10-100bp | 5144 | 2337 | 4998 | 2051 | 5104 | 1817 | 5113 | 1298 | 5073 | 705 |
| 100bp-1Kb | 15690 | 9096 | 15149 | 8268 | 15276 | 7021 | 15309 | 4649 | 15180 | 2754 |
| 1Kb-5Kb | 5073 | 4133 | 4990 | 4062 | 4991 | 3375 | 4993 | 2325 | 4843 | 1438 |
| 5Kb-10Kb | 787 | 839 | 791 | 800 | 801 | 726 | 797 | 521 | 786 | 337 |
| 10Kb-20Kb | 466 | 394 | 472 | 403 | 476 | 384 | 468 | 274 | 439 | 183 |
| 20Kb-30Kb | 155 | 97 | 158 | 110 | 161 | 88 | 163 | 75 | 143 | 44 |
| 30Kb-50Kb | 96 | 87 | 98 | 87 | 99 | 74 | 97 | 56 | 82 | 38 |
| 50Kb-100Kb | 55 | 43 | 57 | 41 | 59 | 41 | 57 | 35 | 50 | 20 |
| **>100Kb** | **20** | **18** | **21** | **17** | **22** | **14** | **25** | **9** | **22** | **5** |
| Total blocks | 27703 | 17140 | 26953 | 15919 | 27210 | 13602 | 27241 | 9285 | 26832 | 5552 |

#### Table S13 Detailed information on introgressed blocks (i-blocks) in m1 and s1 genomes

| **Description** | | **>=2 occurrences of i-allele** | | **>=4 occurrences of i-allele** | | **>=6 occurrences of i-allele** | | **>=8 occurrences of i-allele** | | **=10 occurrences of i-allele** | |
|---|---|---|---|---|---|---|---|---|---|---|---|
| | | **m1 pop** | **s1 pop** | **m1 pop** | **s1 pop** | **m1 pop** | | **m1 pop** | **s1 pop** | **m1 pop** | **s1 pop** |
| **The i-blocks contain >=1 intro sites** | No. of i-blocks | 27703 | 17140 | 26953 | 15919 | 27210 | 13602 | 27241 | 9285 | 26832 | 5552 |
| | No. of scaffolds with i-blocks | 18 | 18 | 18 | 18 | 18 | 18 | 18 | 18 | 18 | 18 |
| | Total length of i-blocks (Mb) | 45.33 | 39.61 | 45.50 | 38.80 | 46.07 | 33.61 | 46.33 | 24.21 | 43.14 | 15.57 |
| | % of the genome | 22.58 | 19.74 | 22.67 | 19.33 | 22.96 | 16.75 | 23.09 | 12.06 | 21.49 | 7.76 |
| **The i-blocks contain >=2 intro sites** | No. of i-blocks | 9742 | 8228 | 9758 | 7896 | 9964 | 6376 | 9963 | 3874 | 9741 | 2219 |
| | No. of scaffolds with i-blocks | 18 | 18 | 18 | 18 | 18 | 18 | 18 | 18 | 18 | 18 |
| | Total length of i-blocks (Mb) | 30.63 | 28.73 | 31.37 | 28.24 | 30.07 | 24.57 | 32.29 | 16.00 | 29.54 | 9.22 |
| | % of the genome | 15.26 | 14.31 | 15.63 | 14.07 | 15.98 | 12.24 | 16.09 | 7.97 | 14.72 | 4.59 |
| **The i-blocks contain >=3 intro sites** | No. of i-blocks | 4805 | 5224 | 4880 | 5106 | 5007 | 4035 | 5004 | 2302 | 4883 | 1302 |
| | No. of scaffolds with i-blocks | 18 | 18 | 18 | 18 | 18 | 18 | 18 | 18 | 18 | 18 |
| | Total length of i-blocks (Mb) | 23.01 | 22.94 | 23.68 | 22.46 | 24.30 | 19.48 | 24.56 | 11.65 | 21.75 | 7.41 |
| | % of the genome | 11.47 | 11.43 | 11.80 | 11.19 | 12.11 | 9.71 | 12.24 | 5.80 | 10.84 | 3.69 |
| **The i-blocks contain >=4 intro sites** | No. of i-blocks | 2861 | 3661 | 2926 | 3628 | 3006 | 2868 | 3001 | 1562 | 2947 | 901 |
| | No. of scaffolds with i-blocks | 18 | 18 | 18 | 18 | 18 | 18 | 18 | 18 | 18 | 18 |
| | Total length of i-blocks (Mb) | 18.60 | 19.00 | 19.19 | 18.55 | 19.75 | 16.53 | 19.71 | 9.43 | 17.21 | 6.03 |
| | % of the genome | 9.27 | 9.47 | 9.56 | 9.24 | 9.84 | 8.23 | 9.82 | 4.70 | 8.58 | 3.01 |
| **The i-blocks contain >=5 intro sites** | No. of i-blocks | 1916 | 2688 | 1955 | 2706 | 2013 | 2159 | 2016 | 1146 | 1981 | 660 |
| | No. of scaffolds with i-blocks | 18 | 18 | 18 | 18 | 18 | 18 | 18 | 18 | 18 | 18 |
| | Total length of i-blocks (Mb) | 16.12 | 15.69 | 16.61 | 15.25 | 17.03 | 13.62 | 17.06 | 7.86 | 14.81 | 4.91 |
| | % of the genome | 8.03 | 7.82 | 8.27 | 7.60 | 8.49 | 6.78 | 8.50 | 3.92 | 7.38 | 2.44 |

#### Table S14 High-confidence non-introgressable j-blocks

| | >=1 j-sites | >=2 j-sites |
|---|---|---|
| No. of j-blocks (No. scaffolds with j-blocks) | 1,189 (184) | 168 (44) |
| Length of j-blocks - Range (mean) | 3 bp-43.51 Kb (1,010 bp) | 23 bp-35.75 Kb (1,062 bp) |
| No. of j-sites in a block – Range (total non-i | 1-6 bp (1,443 bp) | 2-6 bp (422 bp) |
| Total length of j-blocks (% of the genome) | 1,201,823 bp (0.51%) | 178,520 bp (0.075%) |
| No. of genes within j-blocks | 328 | 39 |
| No. of genes containing j-sites | 171 | 19 |

A j-block, unless explicitly stated, should have >= 2 non-introgressable sites (j-sites).

#### Table S15 All functional genes within non-introgressable blocks (j-blocks) between *R. mucronata* and *R. stylosa*

| In *Rhizophora* | | | | In *Arabidopsis thaliana* | |
|---|---|---|---|---|---|
| **Gene** | **j-sites** | **L(aa)** | **sites** | **Gene** | **Function** |
| *RM_76501.12* | 2 | 1,404 | 6 | *AT4G21820* | binding / calmodulin binding protein |
| ***RM_76773.10*** | **3** | **255** | **4** | *AT2G14110* | Haloacid dehalogenase-like hydrolase (HAD) superfamily protein. Participating in pollen germination and tube growth (*47*). |
| *RM_76921.24* | 2 | 539 | 0 | *AT5G04980* | DNAse I-like superfamily protein |
| *RM_76929.4* | 2 | 186 | 3 | *AT4G31940* | CYP82C4 (cytochrome P450, family 82, subfamily C, polypeptide 4). The gene encodes a cytochrome P450 enzyme, CYP82C. It is involved in cellular response to iron ion, oxidation-reduction process, response to iron ion,sideretin [illegible] |
| ***RM_76929.10*** | **2** | **294** | **3** | *AT1G55490* | CPN60B (chaperonin 60 beta). encodes the beta subunit of the chloroplast chaperonin 60, a homologue of bacterial GroEL. Mutants in this gene develops lesions on its leaves, expresses systemic acquired resistance (SAR) and develops accelerated cell death to heat shock stress. Other names: CPN60B, CPN60BETA1, CPNB1, LEN1. This gene can participate in embryo and seed development (*50*, *89*). |
| *RM_76932.33* | 1 | 56 | 0 | unknown | Function unknown. |
| *RM_76963.8* | 2 | 400 | 2 | *AT1G75380* | BBD1 (bifunctional nuclease in basal defense response 1). Involved in defense response to fungus, negative regulation of transcription, DNA-templated,protein ubiquitination, regulation of histone deacetylation. Playing a role in the response to cold stress (*90*). |
| ***RM_76979.9*** | **2** | **199** | **1** | *AT3G15510* | NAC2 (NAC domain containing protein 2). Involved in the regulation of stamen development (*48*), embryonic development (*51*) and stress response (*91*, *92*). |
| *RM_77019.26* | 3 | 351 | 0 | *AT1G29660* | GDSL-like Lipase/Acylhydrolase superfamily protein. Enzyme group with broad substrate specificity that may catalyze acyl-transfer or hydrolase reactions with lipid and non-lipid substrates. Expressed during flower, leaf development and plant embryo globular stage. Expressed in flower, leaf, plant embryo, root, shoot and stem tissues. Participating in both early and late stages of female gametophyte development (*93*). |
| *RM_77067.37* | 2 | 1,903 | 1 | *AT1G09910* | Rhamnogalacturonate lyase family protein. Also known as: F21M12.30; F21M12_30. |

| | | | | | |
|---|---|---|---|---|---|
| ***RM_77078.7*** | ***3*** | **1,415** | ***1*** | *AT5G11530* | EMF1 (embryonic flower 1). Involved in regulating reproductive development (*45*, *46*). |
| *RM_77327.1* | 2 | 585 | 2 | *AT2G39090* | APC7 (tetratricopeptide repeat (TPR)-containing protein). Also known as anaphase-promoting complex 7; AtAPC7; T7F6.26; T7F6_26. Involved in cell cycle, cell division,protein ubiquitination. Located in nucleus. |
| ***RM_77333.68*** | **2** | **755** | **1** | *AT1G08520* | ALB1 (ALBINA 1). Encodes the CHLD subunit of the Mg-chelatase enzyme involved in chlorophyll biosynthesis. Located in chloroplast and extracellular regions. Participating in embryo and seed development (*52*). Lines carrying recessive mutations of this locus are white and seedling lethal. |
| *RM_77333.219* | 2 | 167 | 0 | *AT5G52370* | 28S ribosomal S34 protein. Involved in biological process, response to cold. Located in chloroplast, mitochondrion. Expressed in guard cell. |
| *RM_77333.222* | 2 | 215 | 1 | *AT1G07530* | SCL14 (SCARECROW-like 14). Encodes a member of the GRAS family of transcription factors. The protein interacts with the TGA2 transcription factor and affects the transcription of stress-responsive genes (*94*). The protein is found in the nucleus and is also exported to the cytoplasm. |
| *RM_77333.230* | 2 | 453 | 2 | *AT5G59380* | MBD6 (methyl-CPG-binding domain 6). Protein containing methyl-CpG-binding domain. Has sequence similarity to human MBD proteins. Involved in gene silencing by interacting with RNA binding proteins (*95*). |
| *RM_77333.290* | 5 | 1,056 | 1 | *AT3G45850* | P-loop containing nucleoside triphosphate hydrolases superfamily protein. FUNCTIONS IN: microtubule motor activity, ATP binding; INVOLVED IN: microtubule-based movement. |
| *RM_77333.292* | 2 | 1,398 | 2 | *AT3G45830* | nuclear factor kappa-B-binding-like protein. Function unknown. |
| ***RM_77530.24*** | **2** | **671** | **2** | *AT3G05420* | ACBP4 (acyl-CoA binding protein 4). Acyl-CoA binding protein with high affinity for oleoyl-CoA. Involved in fatty acid transport. Expressed and function in floral lipid metabolism (*96*). Playing combinatory roles in pollen development (*49*) and distinct roles in seed development (*97*). |

The six bolded genes are involved in flower development and/or gamete production and development (see also Table 2).

j-sites: the number of non-introgressable sites within the gene.

L(aa): amino acid sequence length of the gene.

[a] Site: No. of highly differentiated amino acids between *R. mucronata* and *R. stylosa* (see also Fig. S15).

**Supplementary Figures:**

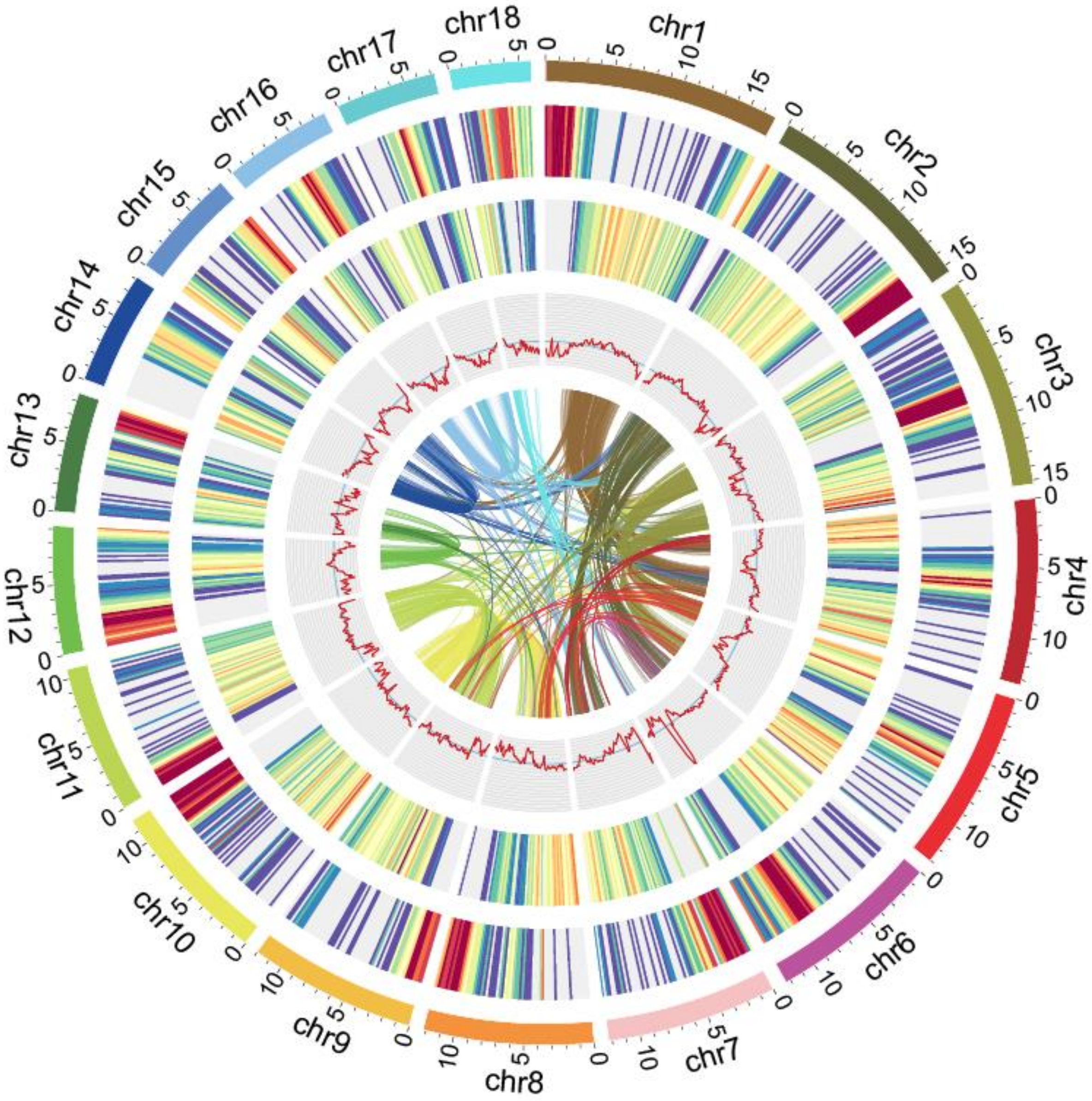


**Figure S1. Features of the *R. mucronata* genome.** Circular tracks represent, from outer to inner, top 18 longest scaffolds (chr1-chr18, with length >= 5Mb), percentage of repeats (2-99%), gene density (0-49), GC content (29.73-51.97%), and the spectrum of collinear analysis (each line connects one pair of homologous genes and a cluster of such lines represents one collinear block). All statistics are calculated in 200 Kb windows.

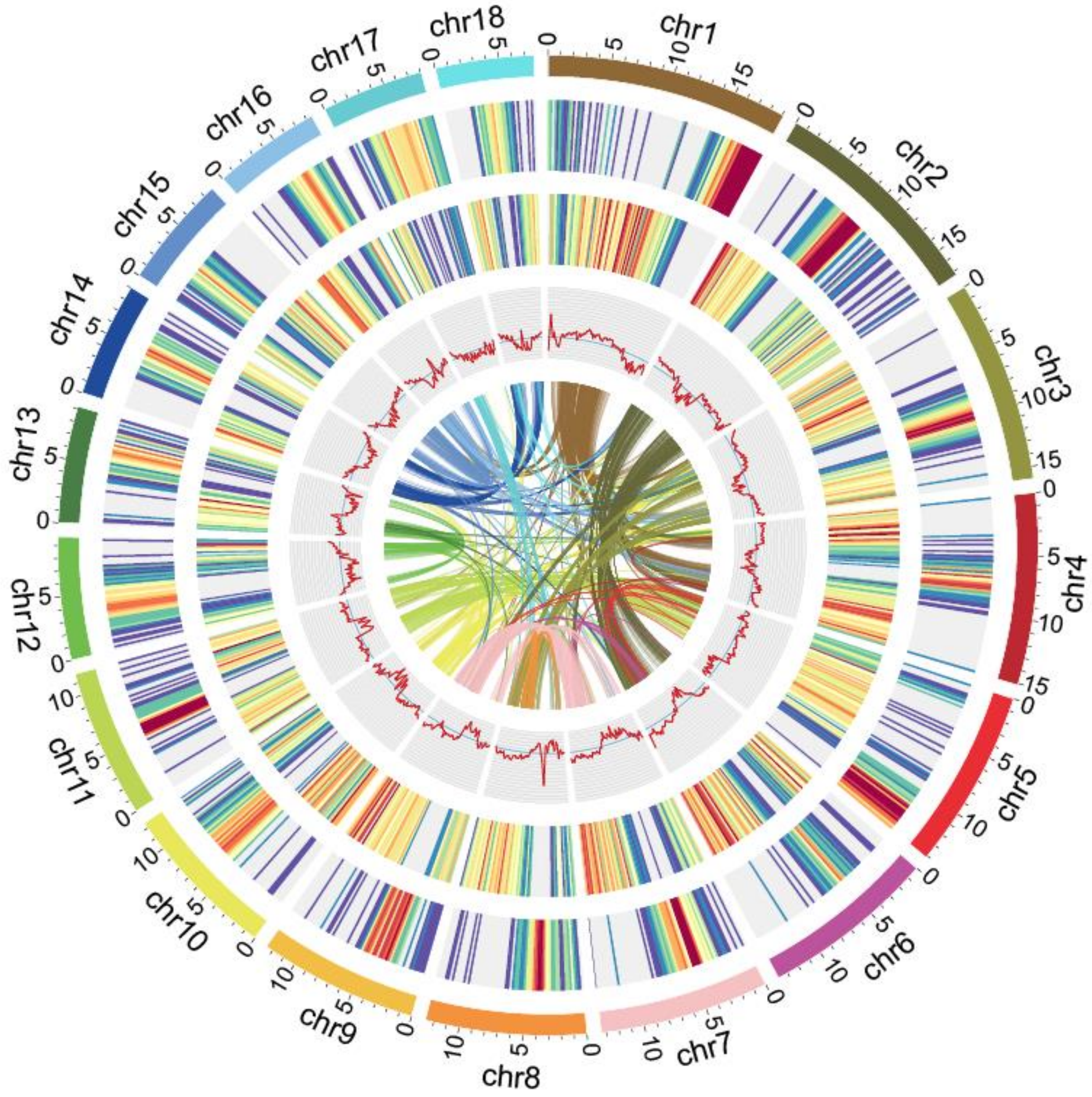


**Figure S2. Features of the *R. stylosa* genome.** Circular tracks represent, from outer to inner, top 18 longest scaffolds (chr1-chr18), percentage of repeats (1-99%), gene density (0-44), GC content (29.61-46.16%), and the spectrum of collinear analysis (each line connects one pair of homologous genes and a cluster of such lines represents one collinear block). All statistics are calculated in 200 Kb windows.

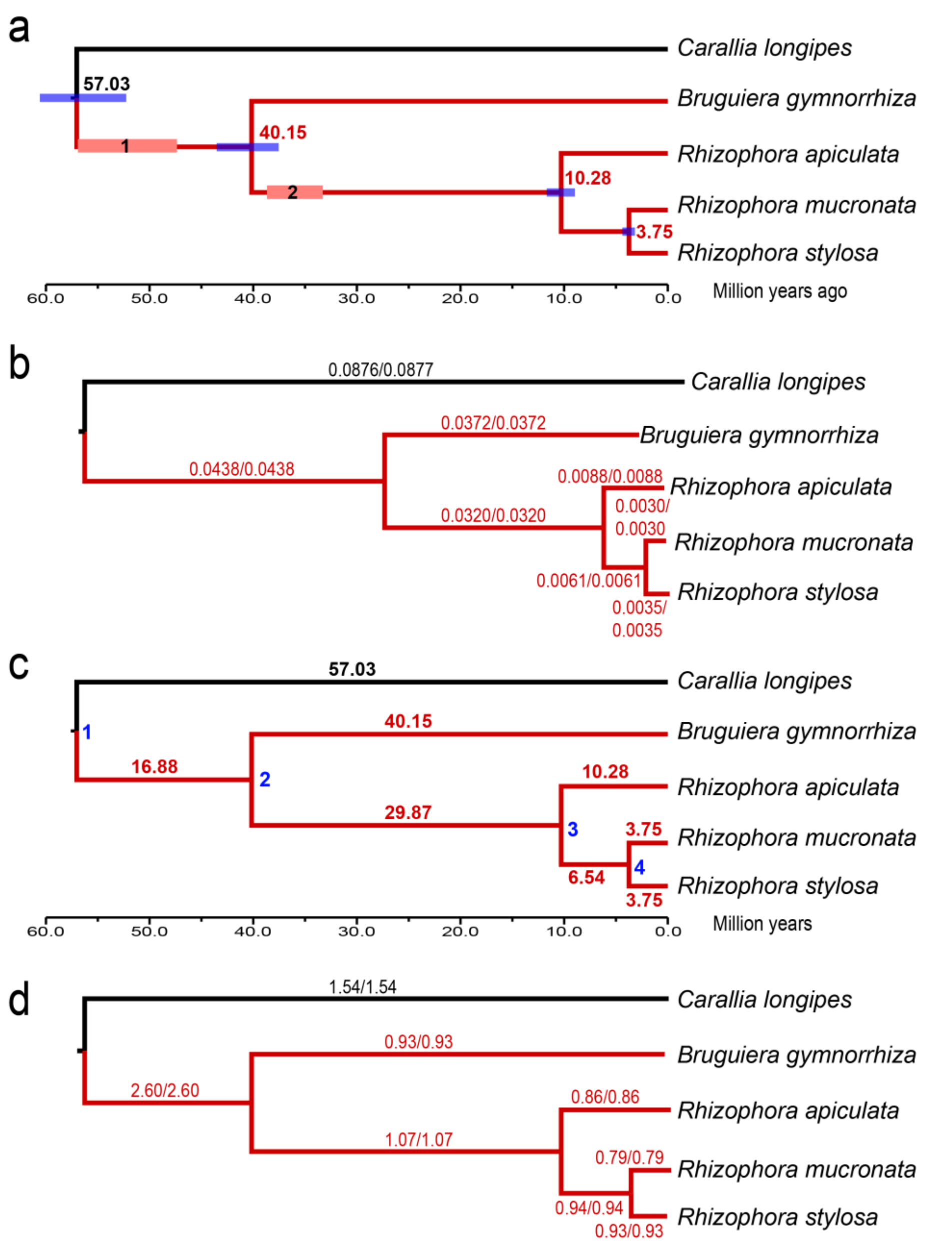


**Figure S3. Phylogenetic relationships and evolution of Rhizophoraceae species.** (a) Phylogenetic relationships and divergence time estimation of five species in the Rhizophoraceae family. The blue bars show 95% credible intervals of divergence time for each node. Red rectangles with numbers represent the earliest known fossil records of mangrove lineages (see Materials and Methods). (b) The phylogenetic tree. The paired numbers above each branch represents branch lengths generated by RAXML (first) and IQTREE (second) using the model GTR+gamma and 1000 bootstrap replicates. All nodes are 100% supported. (c) A Phylogenetic tree showing node numbers and time span of each branch. The nodes are numbered (1-4 in blue) in order to show the divergence time, corresponding to that in the Supplementary Table S3. The time span is above/below each branch, with time unit one million years. (d) Substitution rates (x$10^{-9}$ per site per year) estimated for each branch. The substitution rates are estimated through dividing branch length by the time span of the branch. In all figures, branches in red represent mangrove species.

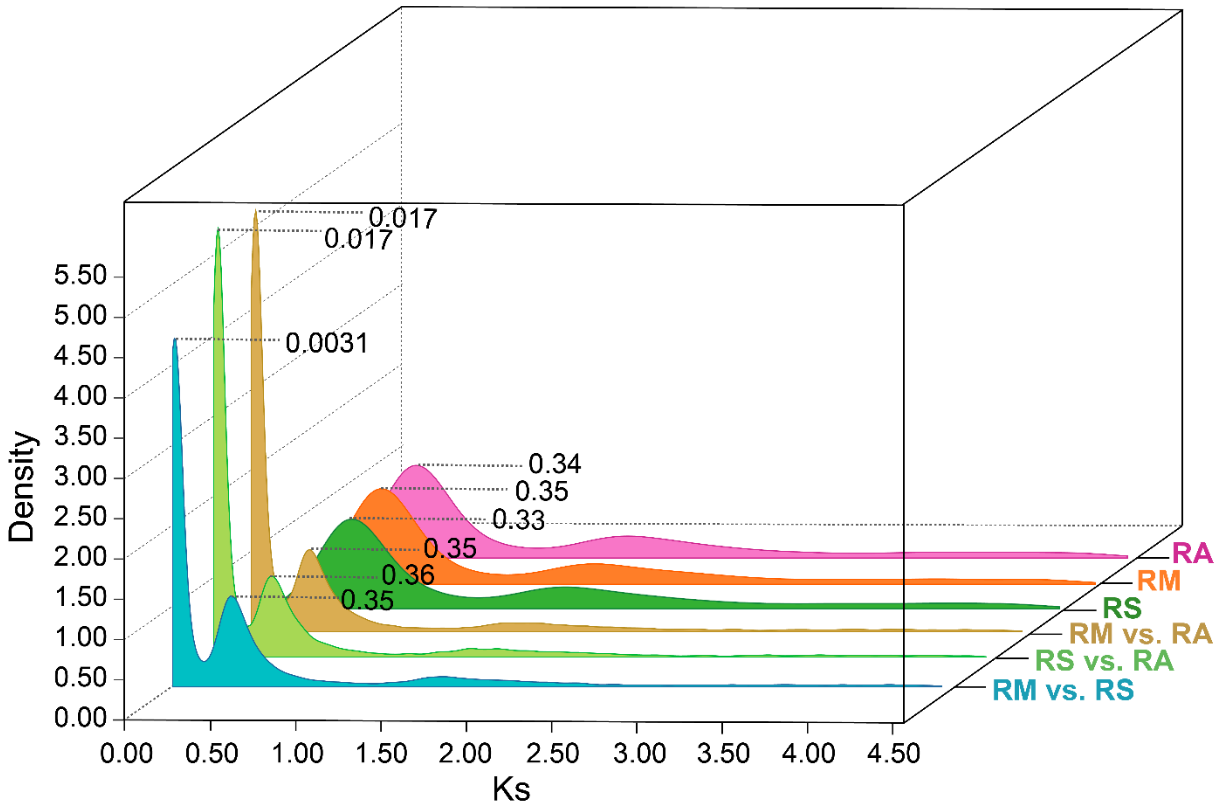


**Figure S4. Intra and inter-specific Ks distributions of three *Rhizophora* species.** Ks values at peaks are next to dotted lines. RM (orange): *R. mucronata*. RS (green): *R. stylosa*. RA (purple): *R. apiculata*. Inter-specific Ks distributions include RM vs. RS (blue), RM vs. RA (light green) and RS vs. RA (dark yellow).

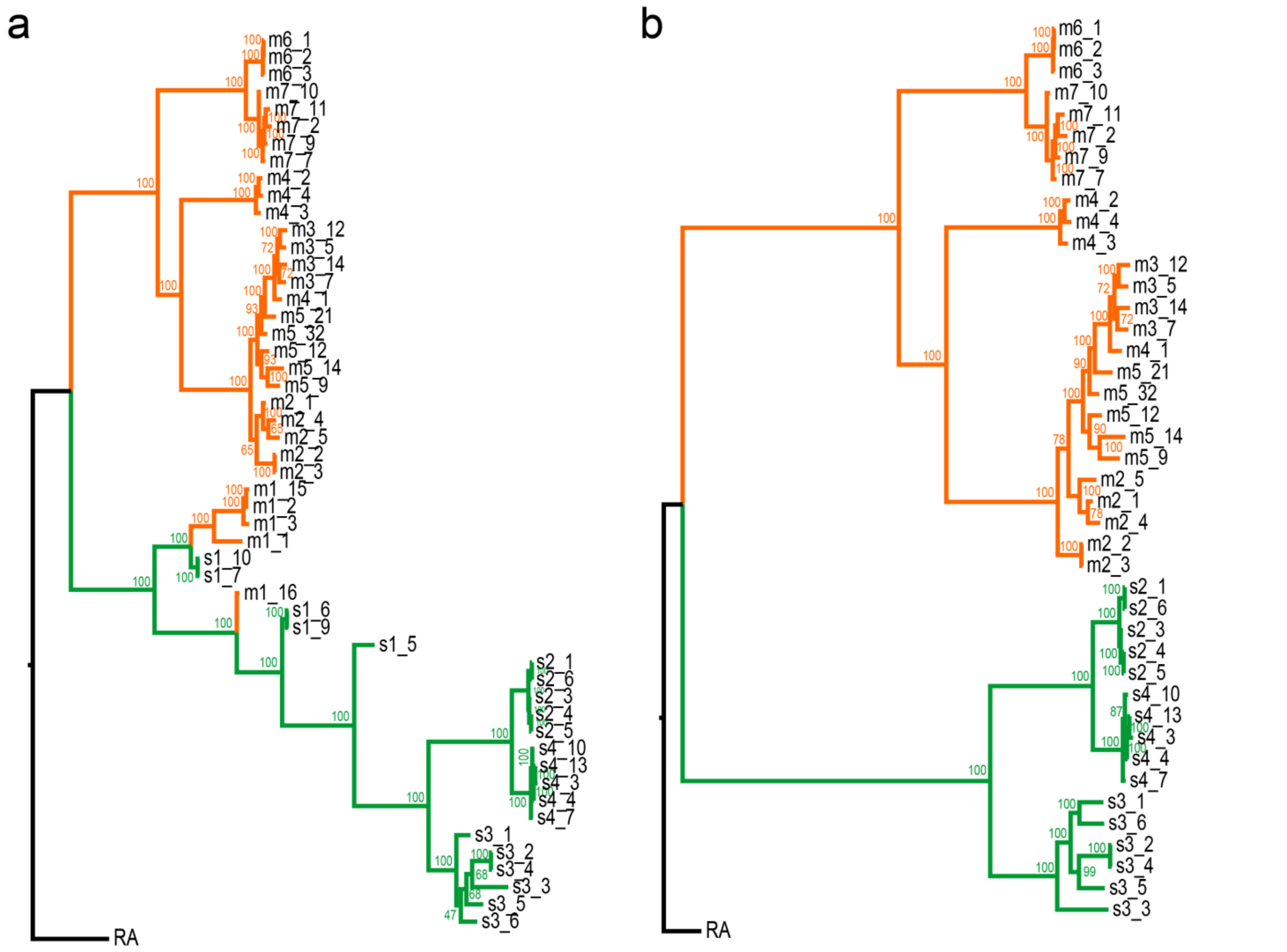


**Figure S5. Phylogenetic relationships of *R. mucronata* and *R. stylosa* samples with (a) or without (b) sympatric populations m1 and s**1. Branches of *R. mucronata* populations (or individuals) are colored in orange while those of *R. stylosa* are in green. The Maximum Likelihood (ML) trees were generated by IQTREE (*31*) with 100 bootstrap replicates. Bootstrap values (supporting rate %) are provided close to each node.

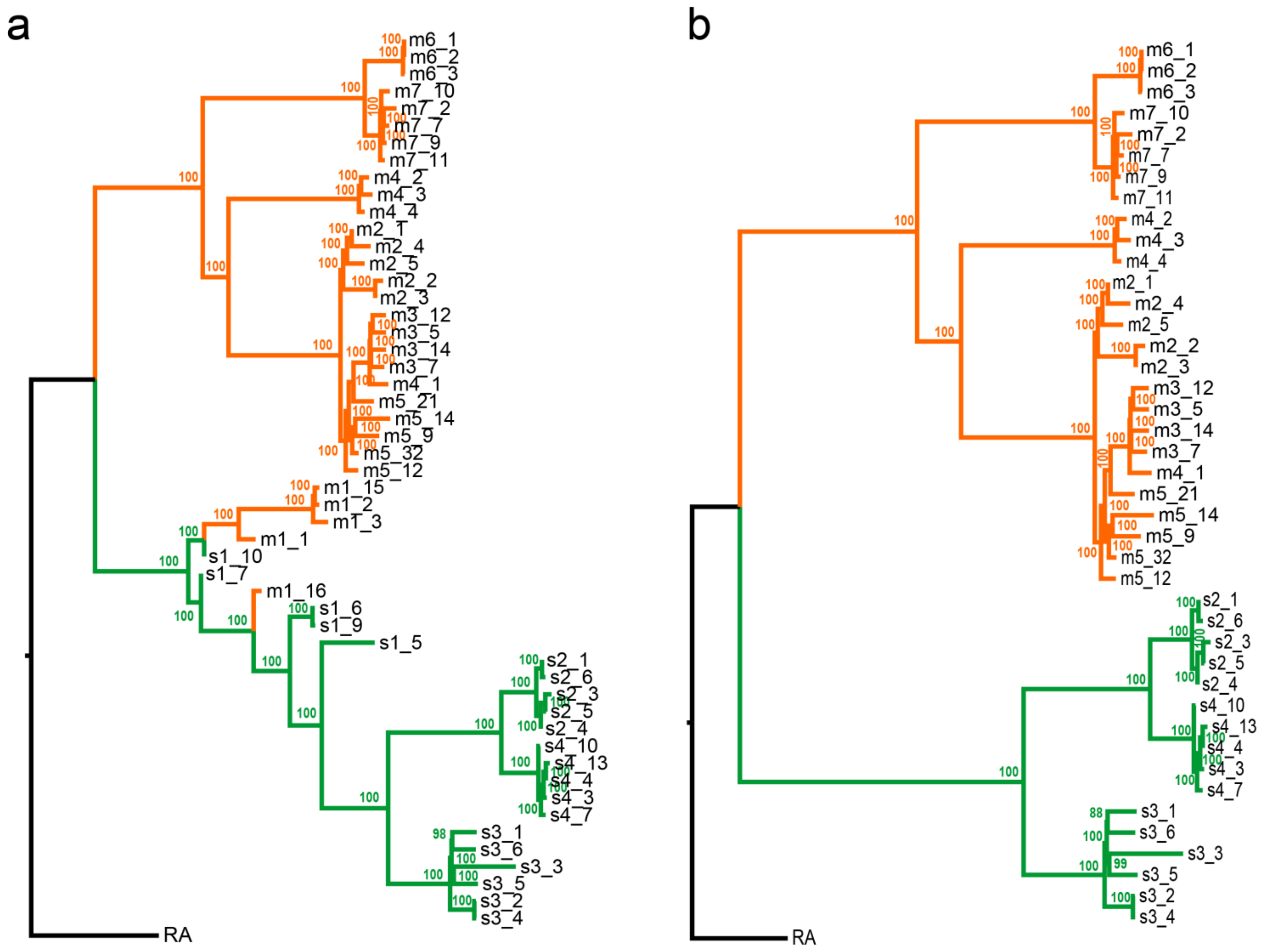


**Figure S6. Phylogenetic relationships of *R. mucronata* and *R. stylosa* samples with (a) or without (b) sympatric populations m1 and s1.** Branches of *R. mucronata* populations (or individuals) are colored in orange while those of *R. stylosa* are in green. The Neighbor-joining (NJ) trees were generated by MEGA7 (*32*) with 100 bootstrap. Bootstrap values (supporting rate %) are provided close to each node.

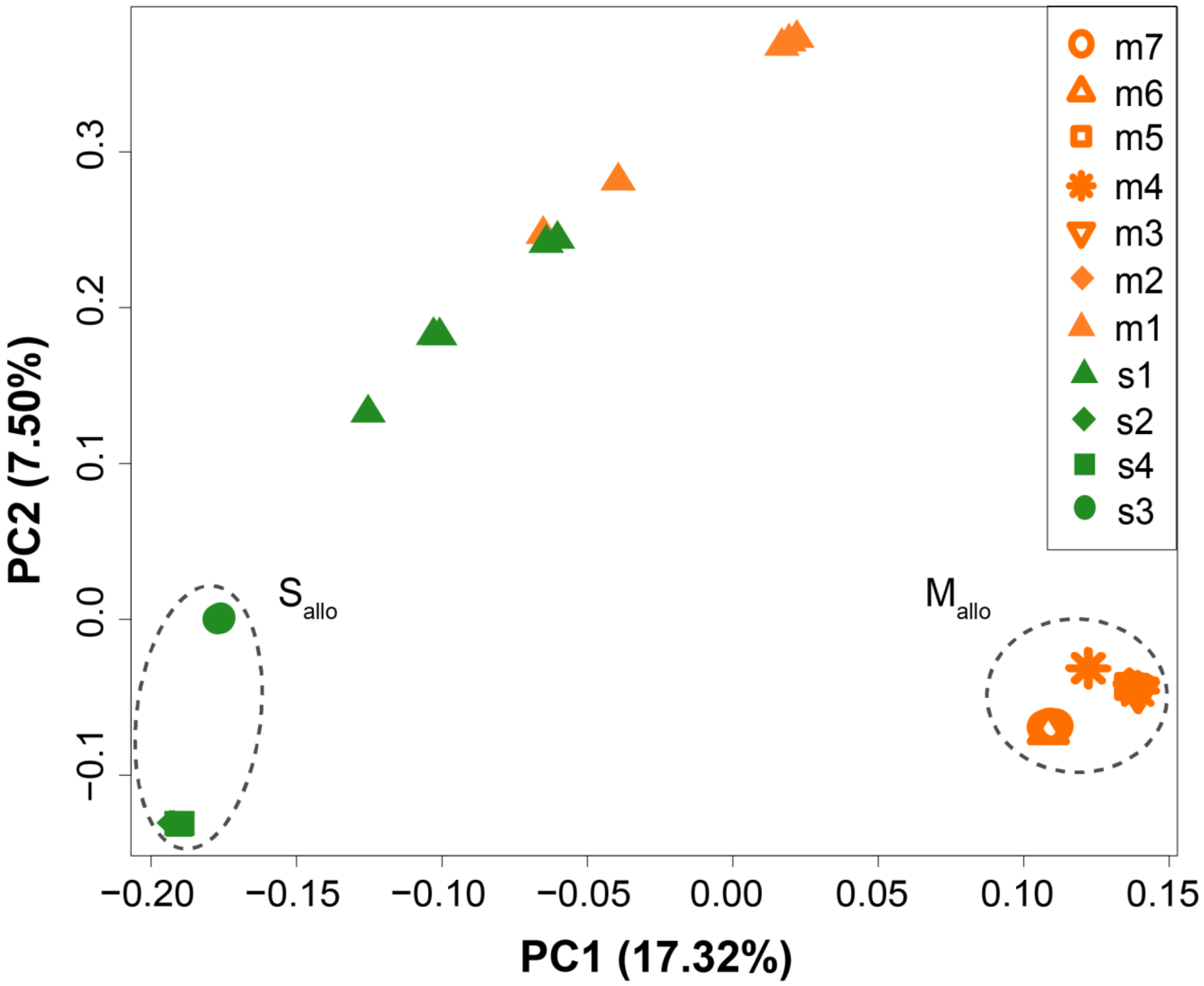


**Figure S7. PCA plot of all populations** (*33*)**.** *R. mucronata* individuals are colored in orange while *R. stylosa* individuals in green. Allopatric populations ($M_{allo}$ and $S_{allo}$) are highlighted by dotted lines. $M_{allo}$ contains populations m2-m7; $S_{allo}$ includes populations s2-s4.

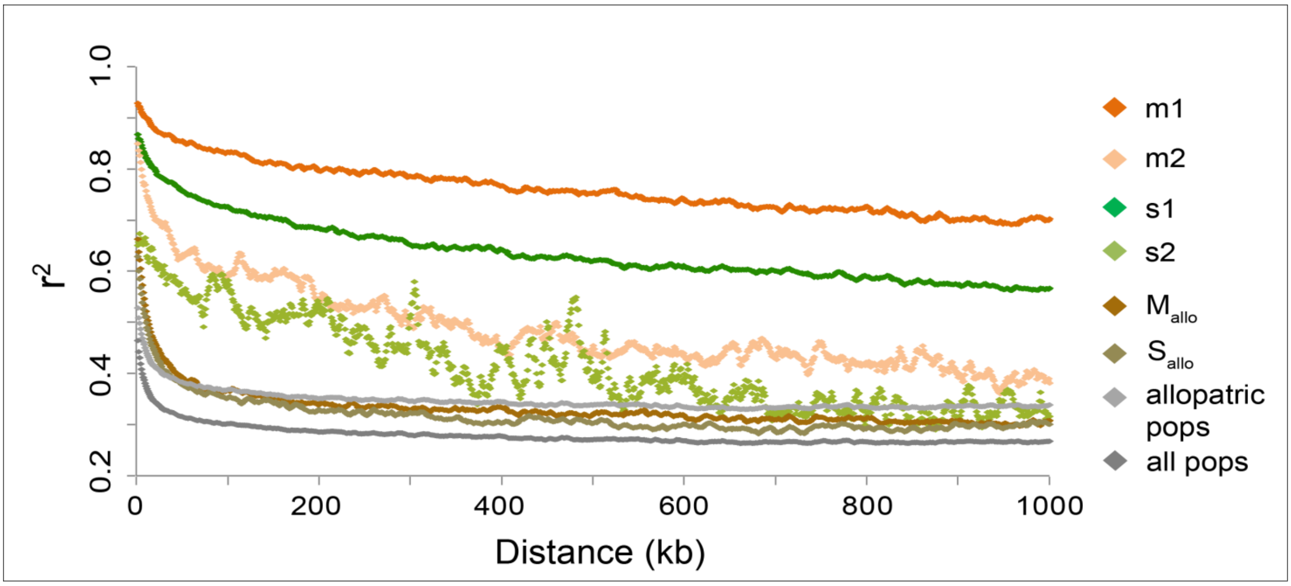


**Figure S8. Decay of linkage disequilibrium in *R. mucronata* and *R. stylosa* populations measured by $r^2$.** $M_{allo}$ contains populations m2-m7; $S_{allo}$ includes populations s2-s4; "allopatric pops" represents all allopatric populations m2-m7 and s2-s4; "all pops" are values for all populations m1-m7 and s1-s4 together.

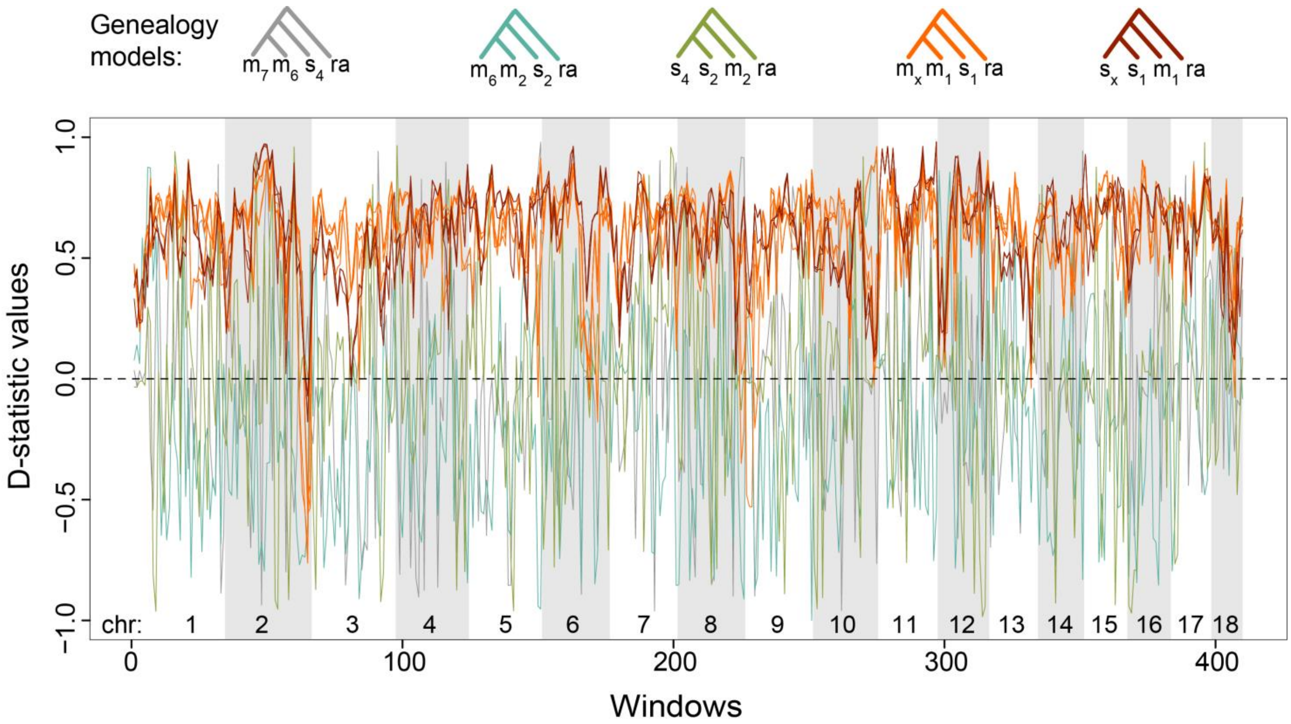


**Figure S9. Patterson's *D* statistic scan across the genome for the genealogy models above, showing genome wide evidence of gene flow between sympatric populations m1 and s1.** The curves are colored by model type. In the last two genealogy models, $m_x$ represents *R. mucronata* population m2, m3, m4, m5, m6 or m7, and $s_x$ represents *R. stylosa* population s2, s3 or s4 (see Table S10 for detail information). The top 18 longest scaffolds (chr1-18) are shown and sibling scaffolds are distinguished by gray shadows.

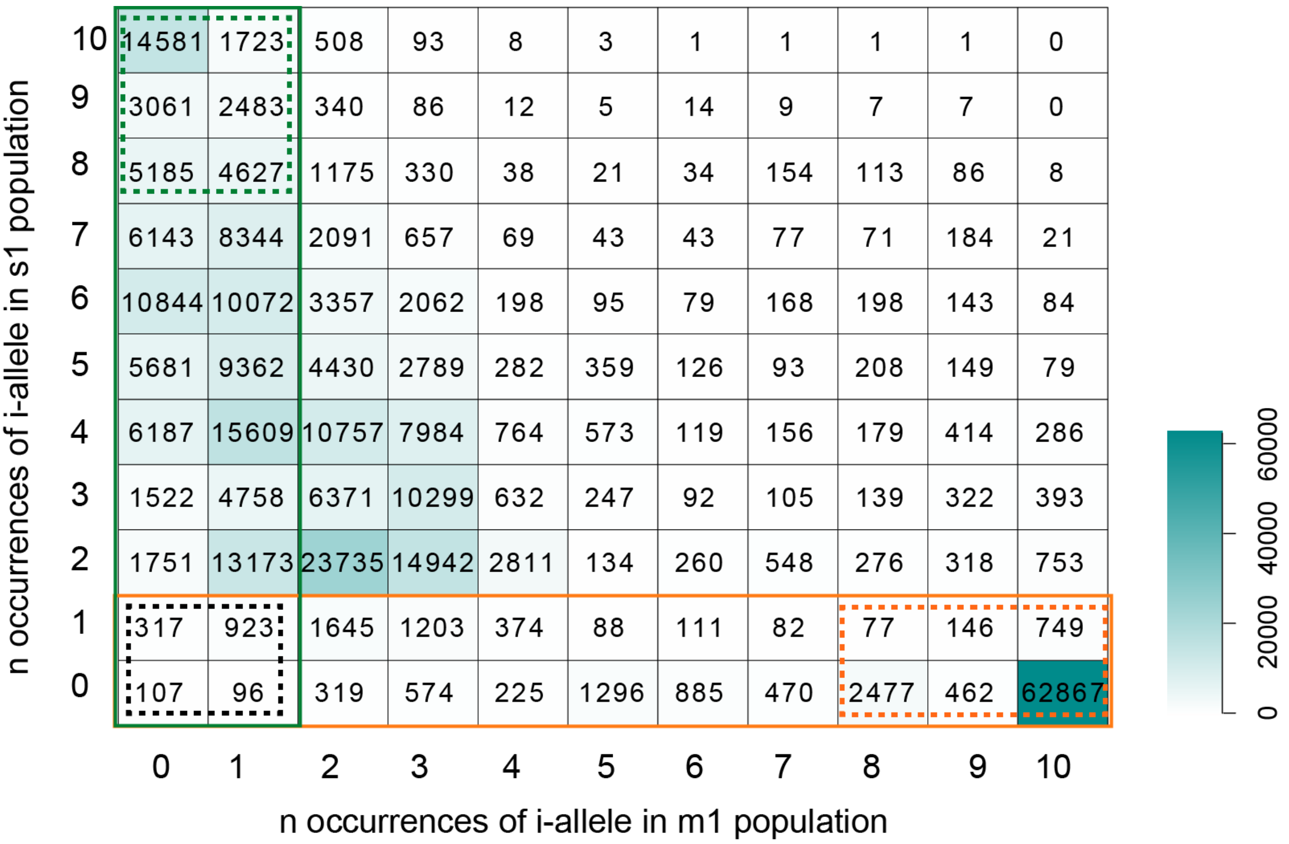


**Figure S10. Distribution of i-allele occurrences in m1 (orange) and s1 (green) populations.** Given the five individuals (or 10 haploid genomes) from each population, the occurrence ranges from 0 to 10. The actual numbers of sites are shown. Sites in the orange (in m1) and green (in s1) solid boxes correspond to the site distributions in Fig. 4B. The orange and green dotted boxes contain the i-sites (with >=8 occurrences of i-allele) in m1 and s1 populations, respectively. The black dotted box shows non-introgressable sites (j-sites) with <=1 occurrences of i-allele both in m1 and s1 samples.

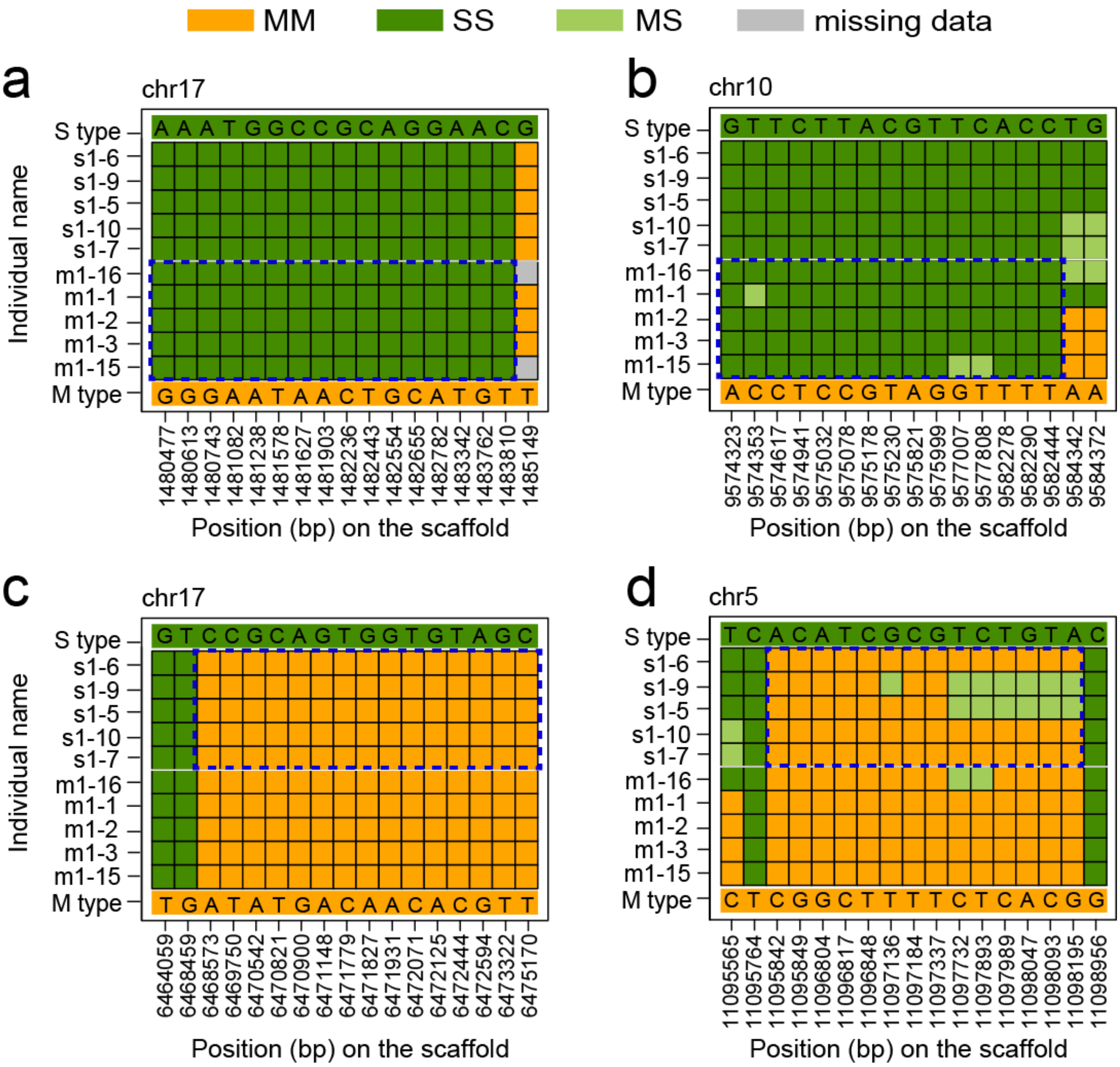


**Figure S11. Examples of i-blocks (in blue dotted boxes) in m1 genomes (a and b) and in s1 genomes (c and d) at the site level.** To make the figures more intuitive and concise, only i-sites and d-sites ($F_{ST}$ > 0.8 between $M_{allo}$ and $S_{allo}$) are shown. Note that i-sites also belong to d-sites (see the text). The top (S type) and bottom (M type) rows in all figures show the dominant bases in $S_{allo}$ and $M_{allo}$, respectively. Each retained row indicates an individual with one vertical line indicating a site. All 10 individuals from the sympatric s1 and m1 populations are shown. Each site is color coded for its genotype: MM (orange), MS (light green) and SS (green) type, where M is for *R. mucronata* and S for *R. stylosa* (see Materials and Methods).

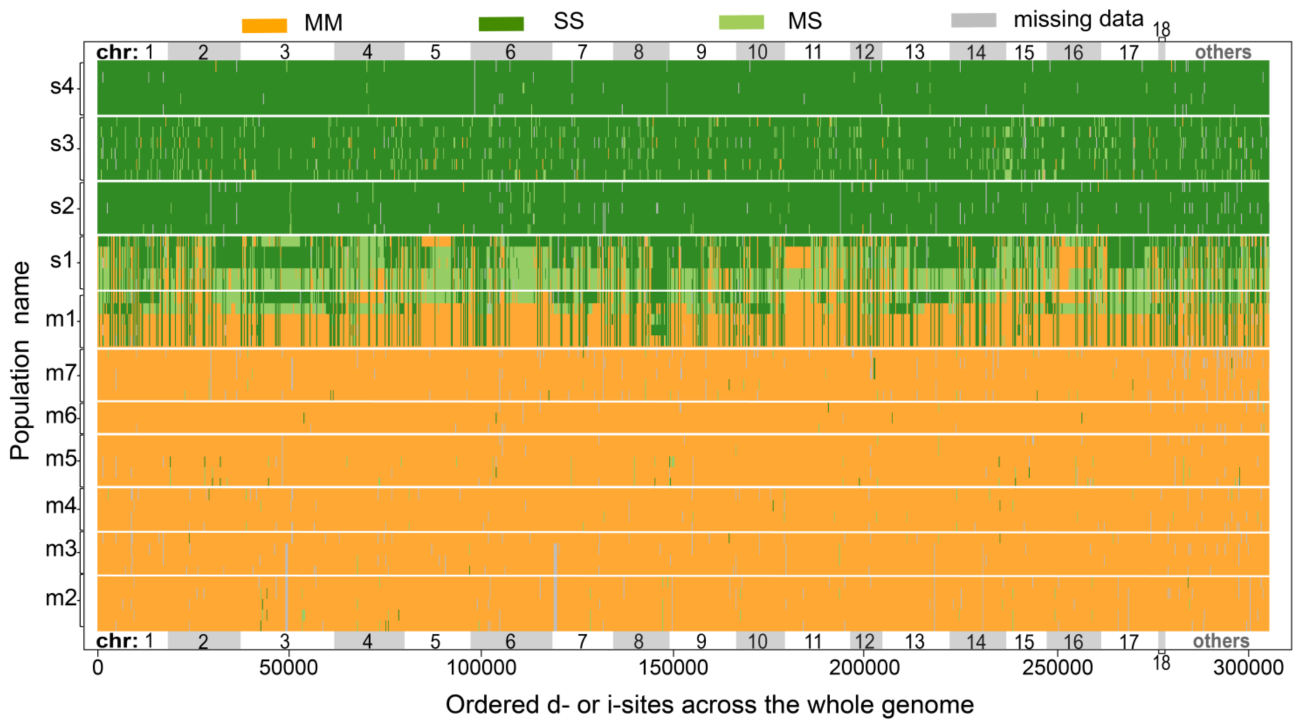


**Figure S12. The genome-wide landscape of i-blocks. All 52 *R. mucronata* and *R. stylosa* individuals are shown.** Top 18 longest scaffolds (chr1-18) and the rest of the genome (others) are marked and sibling scaffolds are distinguished by gray rectangles. In each ideogram, all d- and i-sites are displayed consecutively. Each site is color-coded for the MM, MS, and SS genotypes as in Fig. S11.

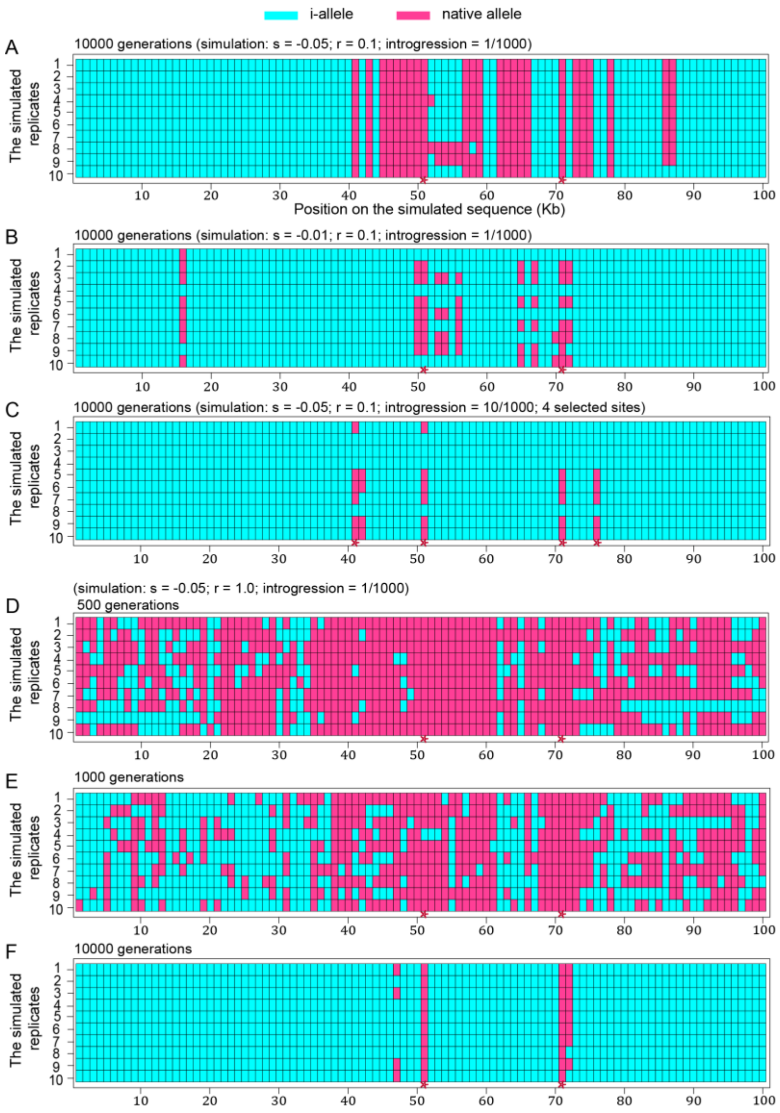


**Figure S13. Simulated introgressions in haploid 100 Kb genomes. Speciation genes (or loci) are marked by red stars at the bottom. Introgressed and non-introgressed (or native) sites are marked in blue and pink, respectively.** (A) Simulated results of 10000 generations under strong selection (s = -0.05), low recombination rate (r = 0.1 for per 100Kb per generation), and low introgression (m=0.001 per generation). Native alleles are not purified at neutral loci. (B) Simulated results of 10000 generations under weak selection (s = -0.01), low recombination rate (r = 0.1 for per 100Kb per generation), and low introgression (m=0.001 per generation). Speciation loci under selection also show introgressions. (C) Simulated results of 10000 generations under strong selection (s = -0.05) plus four loci under selection (#41, #51, #71 and #76), low recombination rate (r = 0.1 for per 100Kb per generation), and high introgression (m=0.01 per generation). Speciation loci under selection have shown introgressions as well. (D-F) Simulated results under strong selection (s = -0.05), high recombination (r = 1.0 for per 100Kb per generation), and low introgression (m=0.001 per generation). Three time points are given. This is closest to the expected pattern.

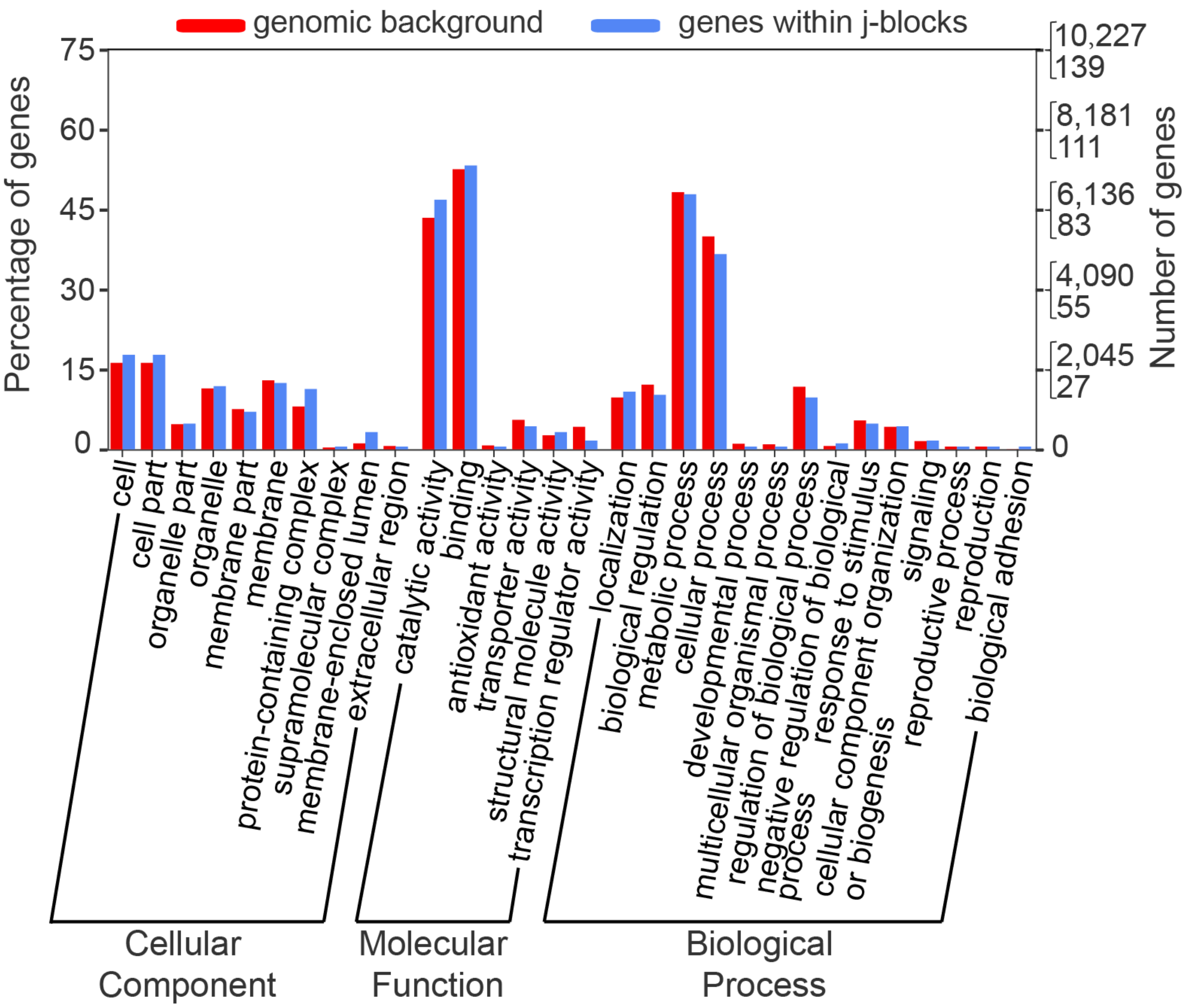


**Figure S14. GO term (level 2) distribution for all genes (328) in the j-blocks (or non-introgressable blocks) between *R. mucronata* and *R. stylosa* (see Table S15).** We use genes from the whole genome as genetic background. In total, 186 genes were assigned to at least one GO term and grouped into three main GO categories and 30 GO terms. WEGO 2.0 (Web Gene Ontology Annotation Plot, http://wego.genomics.org.cn/) (*98*)(<i>98</i>)<sup>98</sup>(J. Ye et al.)<sup>98</sup>(Ye et al., 2018) was used for this analysis.

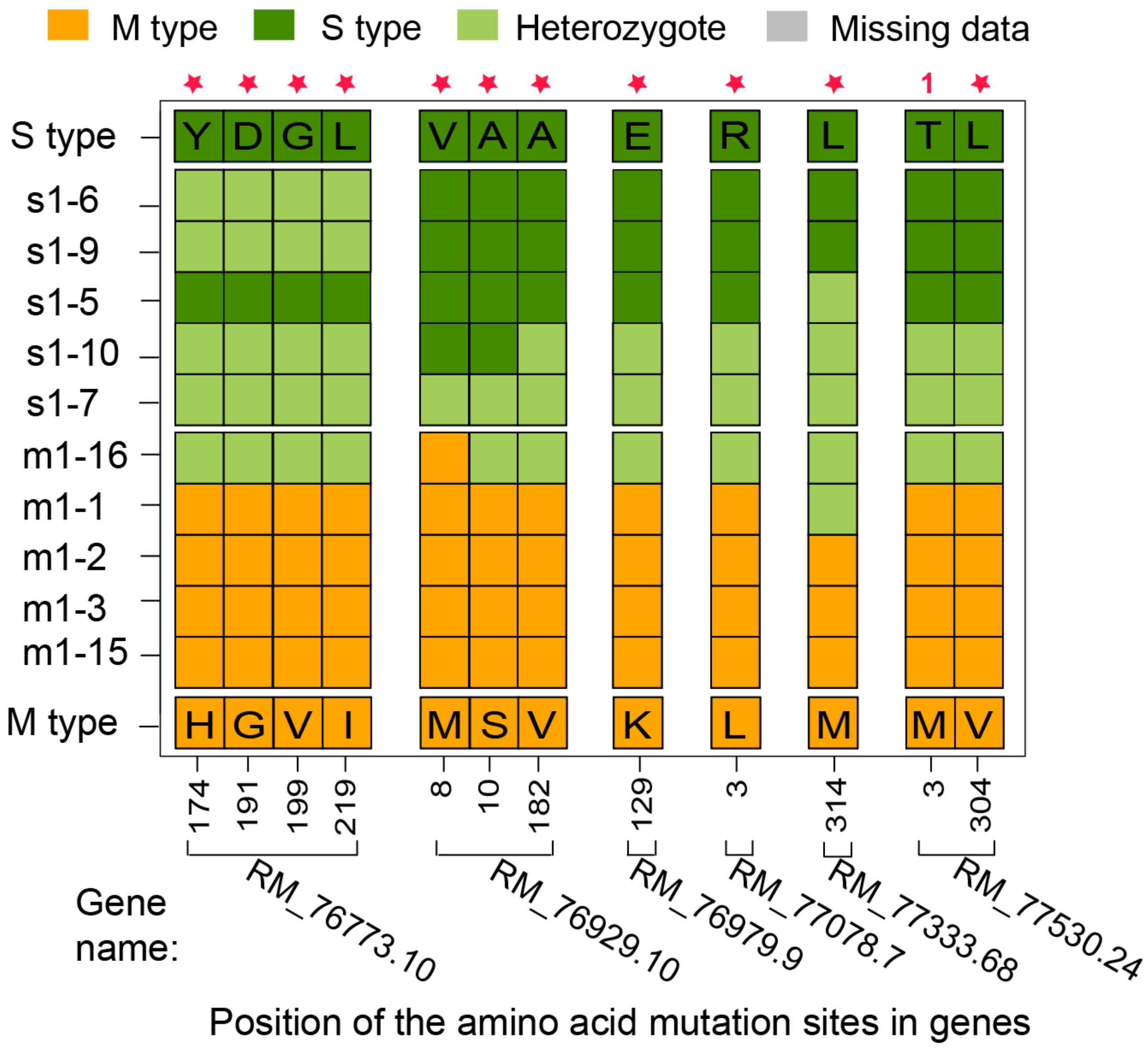


**Figure S15. Highly differentiated amino acids between *R. mucronata* and *R. stylosa* samples in the six genes involved in flower development.** Sites marked by red stars are fixed between allopatric *R. mucronata* (m2-m7) and *R. stylosa* (s2-s4) samples. Only one site contains one (number in red) heterozygote in the s3 (*R. stylosa*) population. Each site is color-coded for the M type (orange), Heterozygote (light green), and S type (green) (M is for allopatric *R. mucronata* dominant amino acid and S for allopatric *R. stylosa* dominant amino acid).

**Supplementary references**


88. N. C. Duke, in *Mangrove Ecosystems: A Global Biogeographic Perspective: Structure, Function, and Services* (2017), Pp.17-53.

89. M. Hajduch *et al.*, Systems Analysis of Seed Filling in Arabidopsis: Using General Linear Modeling to Assess Concordance of Transcript and Protein Expression. *Plant Physiol.* **152**, 2078–2087 (2010).

90. N. Sharma, D. Cram, T. Huebert, N. Zhou, I. A. P. Parkin, Exploiting the wild crucifer Thlaspi arvense to identify conserved and novel genes expressed during a plant's response to cold stress. *Plant Mol. Biol.* **63**, 171–184 (2007).

91. M. Aslam *et al.*, Isolation and characterization of cold responsive NAC gene from Lepidium latifolium. *Mol. Biol. Rep.* **39**, 9629–9638 (2012).

92. T. Chen *et al.*, Effects of tobacco ethylene receptor mutations on receptor kinase activity, plant growth and stress responses. *Plant Cell Physiol.* **50**, 1636–1650 (2009).

93. H. J. Yu, P. Hogan, V. Sundaresan, Analysis of the female gametophyte transcriptome of Arabidopsis by comparative expression profiling. *Plant Physiol.* **139**, 1853–1869 (2005).

94. B. Fode, T. Siemsen, C. Thurow, R. Weigel, C. Gatz, The arabidopsis GRAS protein SCL14 interacts with class II TGA transcription factors and is essential for the activation of stress-inducible promoters. *Plant Cell*. **20**, 3122–3135 (2008).

95. A. P. Parida, A. Sharma, A. K. Sharma, AtMBD6, a methyl CpG binding domain protein, maintains gene silencing in Arabidopsis by interacting with RNA binding proteins. *J. Biosci.* **42**, 57–68 (2017).

96. Z. W. Ye, J. Xu, J. Shi, D. Zhang, M. L. Chye, Kelch-motif containing acyl-CoA binding proteins AtACBP4 and AtACBP5 are differentially expressed and function in floral lipid metabolism. *Plant Mol. Biol.* **93**, 209–225 (2017).

97. A. S. Hsiao *et al.*, Arabidopsis cytosolic acyl-CoA-binding proteins ACBP4, ACBP5 and ACBP6 have overlapping but distinct roles in seed development. *Biosci. Rep.* **34**, 865–877 (2014).

98. J. Ye *et al.*, WEGO 2.0: A web tool for analyzing and plotting GO annotations, 2018 update. *Nucleic Acids Res.* **46**, W71–W75 (2018).
